# High-Fidelity, Hyper-Accurate, and Evolved Mutants Rewire Atomic Level Communication in CRISPR-Cas9

**DOI:** 10.1101/2023.08.25.554853

**Authors:** Erin Skeens, Souvik Sinha, Mohd Ahsan, Alexandra M. D’Ordine, Gerwald Jogl, Giulia Palermo, George P. Lisi

## Abstract

The Cas9-HF1, HypaCas9, and evoCas9 variants of the Cas9 endonuclease are critical tools to mitigate off-target effects in the application of CRISPR-Cas9 technology. The mechanisms by which mutations in the Rec3 domain mediate specificity in these variants are poorly understood. Here, solution NMR and molecular dynamics simulations establish the structural and dynamic effects of high-specificity mutations in Rec3, and how they propagate the allosteric signal of Cas9. We reveal conserved structural changes and peculiar dynamic differences at regions of Rec3 that interface with the RNA:DNA hybrid, transducing chemical signals from Rec3 to the catalytic HNH domain. The variants remodel the communication sourcing from the Rec3 α-helix 37, previously shown to sense target DNA complementarity, either directly or allosterically. This mechanism increases communication between the DNA mismatch recognition helix and the HNH active site, shedding light on the structure and dynamics underlying Cas9 specificity and providing insight for future engineering principles.

---

CRISPR-Cas9 is a revolutionary genome editing system widely adapted for bioengineering purposes^1^. Its use as a therapeutic for human diseases, often regarded as the pinnacle of Cas9 application, is hampered by the occurrence of off-target effects^2^. To overcome this limitation, directed evolution and extensive engineering of the Cas9 enzyme have been performed, leading to a set of promising variants that exhibit enhanced specificity toward target DNA^3–5^. Interestingly, many of these variants include mutations distal from the catalytic sites, though the mechanisms that confer specificity from long-range mutational effects remain unclear. Elucidation of these specificity-enhancing mechanisms is critical for the rational design and further improvement of Cas9 variants that mitigate off-target cleavage in biomedical applications.

The *Streptococcus pyogenes* Cas9 (SpCas9), the most broadly utilized Cas9 enzyme, has a multi-domain architecture consisting of two nuclease domains, HNH and RuvC, a Protospacer Adjacent Motif (PAM)-interacting domain (PI), and a recognition lobe that mediates nucleic acid binding through three distinct subdomains, Rec1–3 (Fig. 1a)^6, 7^. Upon recognition of the short PAM sequence, Cas9 locally unwinds the double-stranded DNA (dsDNA) and a Cas9-bound guide RNA (gRNA) forms an RNA:DNA hybrid with the target DNA strand (TS)^8^. Coordinated cleavage of the TS and non-target strand (NTS) then occurs via the HNH and RuvC nucleases, respectively. Biophysical studies and molecular dynamics (MD) simulations have indicated that the molecular function of Cas9 is governed by an intricate allosteric response driven by intrinsic dynamics, controlling the activation of the catalytic HNH domain^9,10^. Indeed, the flexibility of HNH ensures DNA cleavage at the proper location^11, 12^, but its conformational dynamics were shown to be dependent on the motions of the spatially distant Rec3 domain^3, 9, 13, 14^, which was therefore regarded as an “allosteric effector” of HNH function^3^. Nevertheless, the molecular details of this Rec3-mediated activation are unknown.

**Fig. 1.**
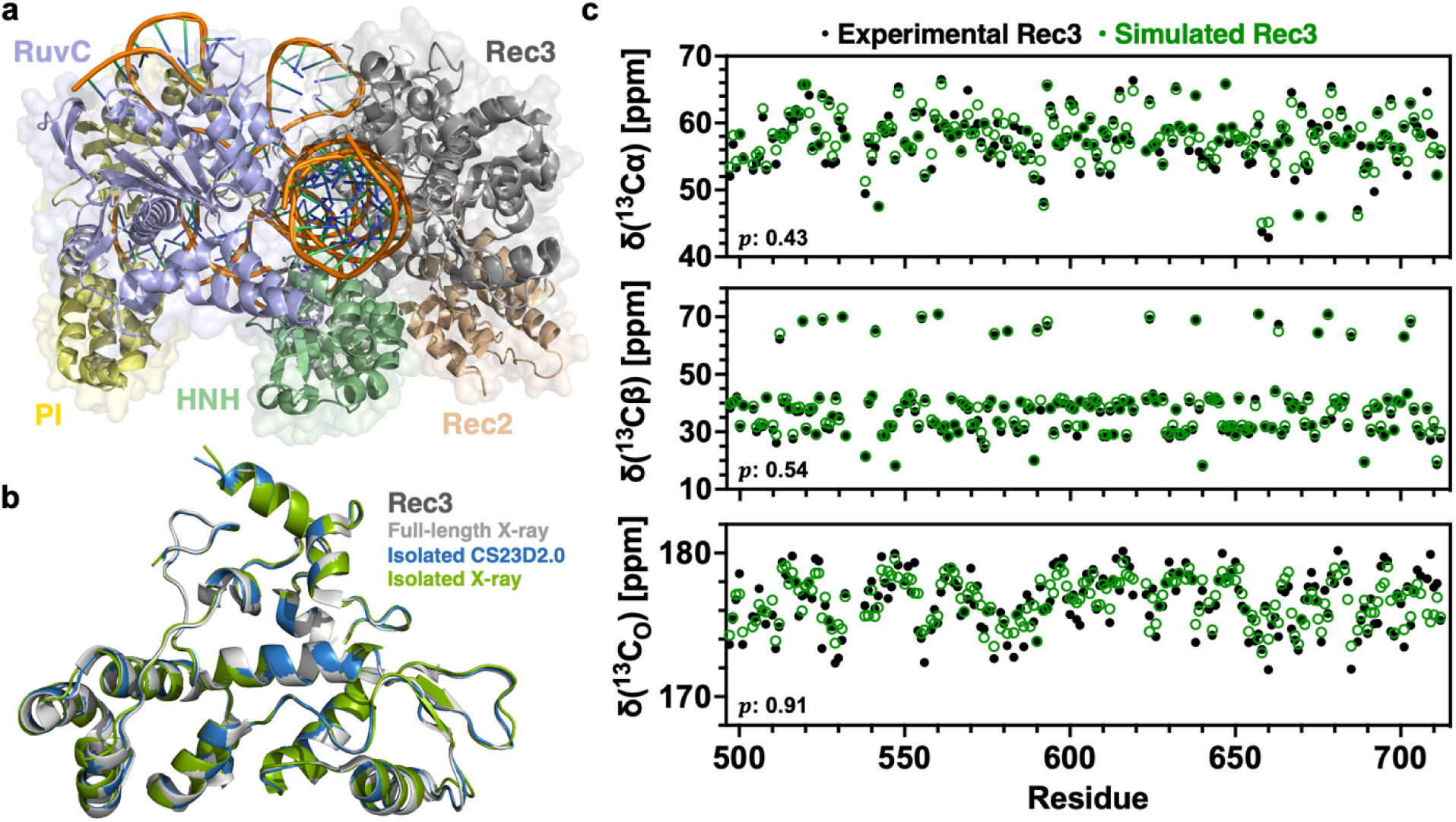
The isolated Rec3 domain structure mirrors Rec3 from full-length Cas9. **a.** Domain arrangement of the *Streptococcus pyogenes* Cas9, highlighting the RuvC (blue), PAM-interacting (PI; yellow), HNH (green), Rec3 (grey), and Rec2 (tan) domains, with the RNA and DNA displayed in orange. **b.** The overlaid structures of Rec3 from the full-length Cas9 X-ray crystal structure (PDB: 4UN3. grey)^7^, our NMR-derived CS23D2.0 model (blue), and the isolated Rec3 X-ray crystal structure (PDB: 8SCA, green), solved here at 1.67 Å, reveal remarkable structural consistency (i.e., Cα RMSD of 0.649 Å between Rec3 in PDB: 8SCA and in PDB: 4UN3; Cα RMSD of 0.598 Å between Rec3 in PDB: 8SCA and in the NMR-predicted model). **c.** Experimental (black dots) and simulated (green dots) chemical shifts of ^13^C_α_ (top), ^13^C_β_ (centre), and ^13^C_O_ (bottom) are plotted for residues in the Rec3 domain. For each atom type, the estimated -values are in the 95% confidence interval, confirming the significance of the overlap between experimental and simulated chemical shift measurements.

Intriguingly, efforts to engineer SpCas9 variants for enhanced specificity revealed that Rec3 is an important driver of off-target recognition. Many notable variants house the majority of their specificity-conferring mutations within the Rec3 domain, including the “high fidelity” Cas9-HF1^4^, the “hyper-accurate” HypaCas9^3^, and the “evolved” evoCas9^5^ variants, suggesting that Rec3 plays a critical role in proofreading the DNA. However, the mechanisms by which mutations in Rec3 mediate the specificity of Cas9, especially considering their distance from the catalytic sites, are poorly understood. Single-molecule experiments have revealed that the HF1 and Hypa mutations alter the conformational equilibrium that controls Cas9 activation^3^ and that high-specificity mutations in Rec3 affect DNA unwinding and the RNA:DNA heteroduplexation process^15–17^. Further atomic-level description of the mutation-induced Cas9 structure and dynamics at the level of Rec3 could contribute to discerning the specificity-enhancing molecular mechanism and the allosteric role of Rec3.

To understand the allosteric contributions of Rec3 to Cas9 specificity and function, we combined experimental and computational techniques to comprehensively characterize the structural and dynamic effects of high-specificity mutations on the Rec3 domain and more broadly, Cas9. We engineered a construct of the isolated Rec3 domain to experimentally probe its biophysical properties and the effect of the HF1, Hypa, and Evo mutations with atomic resolution through solution NMR. In parallel, multi-microsecond (μs) MD simulations of the full-length CRISPR-Cas9 system and its variants were used to describe how the allosteric signal propagates from Rec3 through the full complex. Our findings reveal conserved structural changes and peculiar dynamics within the high-specificity Rec3 variants, which rewires allosteric signalling in CRISPR-Cas9 and increase the communication between the DNA recognition region and the HNH catalytic core.

## RESULTS

### Structure and dynamics of the allosteric Rec3 domain

To probe the contribution of Rec3 to Cas9 function, X-ray crystallography and solution NMR were used to characterize the structure and dynamics of the isolated Rec3 domain, while MD simulations of the full-length CRISPR-Cas9 system were employed to study Rec3 within the larger complex. The combination of these approaches can comprehensively describe allosteric mechanisms in large biomolecules^18–23^. Indeed, solution NMR can assess the domain-specific dynamics with experimental accuracy, while MD simulations can interpret the dynamics in the context of the full-length assembly to understand the allosteric crosstalk and track the signal transmission.

For experimental studies of the Rec3 domain, an isolated construct of Rec3 (residues 497-713) was engineered. A structural model of the isolated Rec3 domain was predicted from the assigned ^1^H-^15^N HSQC NMR spectrum and carbon (Cα, Cβ, C_O_) chemical shifts using the CS23D2.0 server. Fig. 1b shows an overlay of the CS23D2.0 model of the isolated Rec3 domain (blue) with Rec3 from the full-length Cas9 crystal structure (grey; PDB: 4UN3). The NMR-derived CS23D2.0 model is remarkably consistent with Rec3 from the Cas9 crystal structure. Secondary structure analysis of the isolated Rec3 domain from Cα/Cβ chemical shifts and circular dichroism (CD) spectroscopy supports a predominantly alpha-helical structure (Supplementary Fig. 1a,b), providing further indication that the engineered construct maintains the expected fold of the Rec3 domain within the full-length Cas9 protein.

We then determined the structure of the isolated Rec3 domain via X-ray crystallography (PDB: 8SCA at 1.67 Å resolution, Fig. 1b), which also aligns remarkably well with that of Rec3 from the Cas9 crystal structure and the NMR-predicted model, with a Cα root-mean-square deviation (RMSD) of 0.649 Å and 0.598 Å, respectively. There are no major structural differences observed and minor differences are localized to flexible loops. This further confirms the strong NMR-predicted agreement between the isolated Rec3 and the domain within the full-length system. Multi-microsecond (μs) simulations were then performed on a full-length Cas9 complex, where an overall ensemble of ∼16 μs (arising from four replicas of ∼4 μs each) was used to compute Cα, Cβ, and C_O_ chemical shifts of Rec3 with SHIFTX2^24^ for comparison to those measured experimentally by NMR. We detected excellent agreement between the experimental and simulated values (Fig. 1c), which further supports that the isolated Rec3 construct and resulting NMR spectra are valuable for studying the structural and dynamic determinants of Rec3-mediated allostery and specificity (Supplementary Fig. 2). In turn, MD simulations of the full-length Cas9 properly represent the NMR data on the isolated Rec3, which offers the opportunity to study how allosteric information propagates from Rec3 to the rest of the system.

NMR relaxation experiments were conducted to probe the dynamics of Rec3 on timescales relevant to the transmission of allosteric signals. Fast molecular motions on the ps – ns timescale were assessed via order parameters (*S*^2^), calculated from the longitudinal (*R*_1_) and transverse (*R*_2_) relaxation rates and ^1^H-[^15^N] heteronuclear NOE (Figs. 2a,c and Supplementary Fig. 3)^18, 25, 26^. For quantification of slower dynamic processes on the μs – ms timescale, which are classically associated with the transmission of allosteric effects in many enzymes^27, 28^, including the HNH nuclease of Cas9^10^, Carr−Purcell−Meiboom−Gill (CPMG) relaxation dispersion experiments were performed (Fig. 2b,c and Table S1). Residues in Rec3 with motions on the ps – ns timescale are observed in intrinsically dynamic loops and on the face of the domain that does not interact with the RNA:DNA hybrid. Conversely, sites that exhibit μs – ms motions are generally localized to the RNA:DNA hybrid interface, with exchange contributions to relaxation (*R_ex_*) > 4, suggesting that the intrinsic flexibility of the Rec3 domain on the μs – ms timescale is important for facilitating its interaction with the RNA:DNA hybrid.

**Fig. 2.**
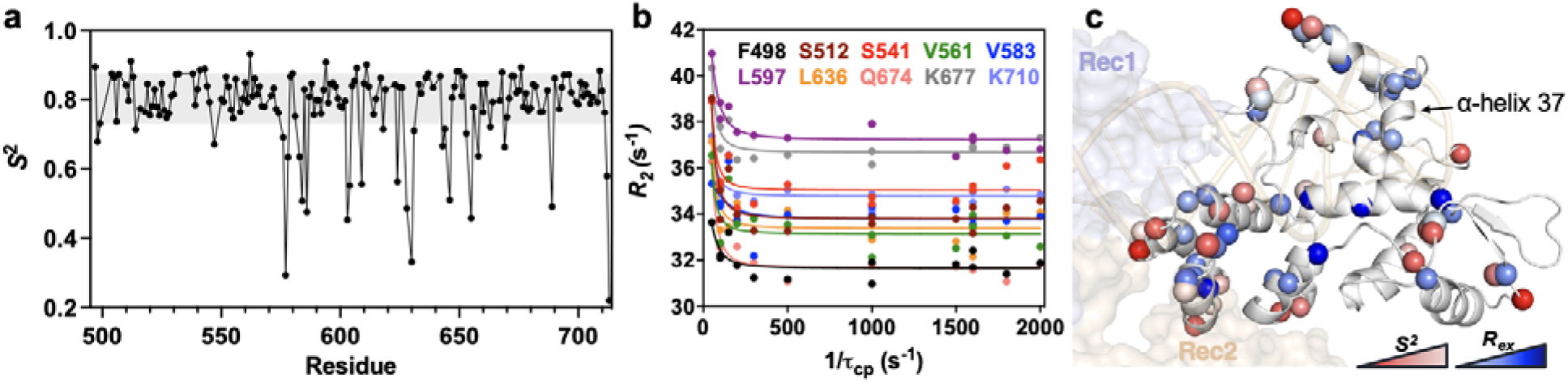
Flexibility of the wild-type (WT) Rec3 domain. **a.** Order parameters (*S^2^*) of WT Rec3 reveal ps – ns timescale motions. The grey shaded area denotes ±1.5σ from the 10% trimmed mean of the data, where order parameters below this area represent residues with ps – ns motions. **b.** A selection of residues in Rec3 exhibiting μs – ms timescale flexibility via representative CPMG relaxation dispersion profiles. **c.** Residues displaying ps – ns motions via *S^2^* and CPMG-detected μs – ms motions are mapped onto the Rec3 structure, shown to display the face of the domain that interacts with the RNA:DNA hybrid. Red spheres represent ps – ns motions (*S^2^*), with a gradient denoting low *S^2^* values as dark red, signifying greater flexibility. Blue spheres denote μs – ms motions (CPMG), with a blue gradient that becomes darker as the exchange contribution to relaxation (*R_ex_*) increases. Rec1, Rec2, and the RNA:DNA hybrid are shown for reference.

### Evidence for a universal structural change in the high-specificity Rec3

To probe whether structural and dynamic changes contribute to the enhanced specificity of the Cas9-HF1, HypaCas9, and evoCas9 variants, we introduced the specificity-conferring mutations into the isolated Rec3 domain to generate HF1 Rec3 (N497A/R661A/Q695A), Hypa Rec3 (N692A/M694A/Q695A/H698A), and Evo Rec3 (Y515N/K526E/R661Q).

A ^1^H-^15^N HSQC spectral overlay of the Rec3 variants with wild-type (WT) Rec3 shows widespread chemical shift perturbations, indicative of local structural changes induced by the mutations (Supplementary Fig. 4), and line-broadened resonances, suggestive of a dynamic change. Analysis of the chemical shift perturbations, deemed statistically significant when 1.5σ above the 10% trimmed mean of all data sets, reveal a highly consistent pattern exhibited by the Rec3 variants (Fig. 3a). Three notable regions spanning residues 510-555, 576-595, and 678-698 are significantly perturbed and exhibit line broadening in all Rec3 variants, despite the disparate locations of the mutations within each construct. Residues exhibiting significant chemical shift perturbations in any single variant, any two variants, and all three variants are mapped onto the Rec3 structure in Fig. 3b, denoted by yellow, orange, and red spheres, respectively, to highlight residues that are most frequently perturbed by the mutations. The perturbations are largely present on the face of the Rec3 structure that interfaces with the RNA:DNA hybrid at the PAM distal ends. Non-specific perturbations observed in only one of the three variants (Fig. 3b, yellow spheres) occur predominantly around the sites of mutation.

**Fig. 3.**
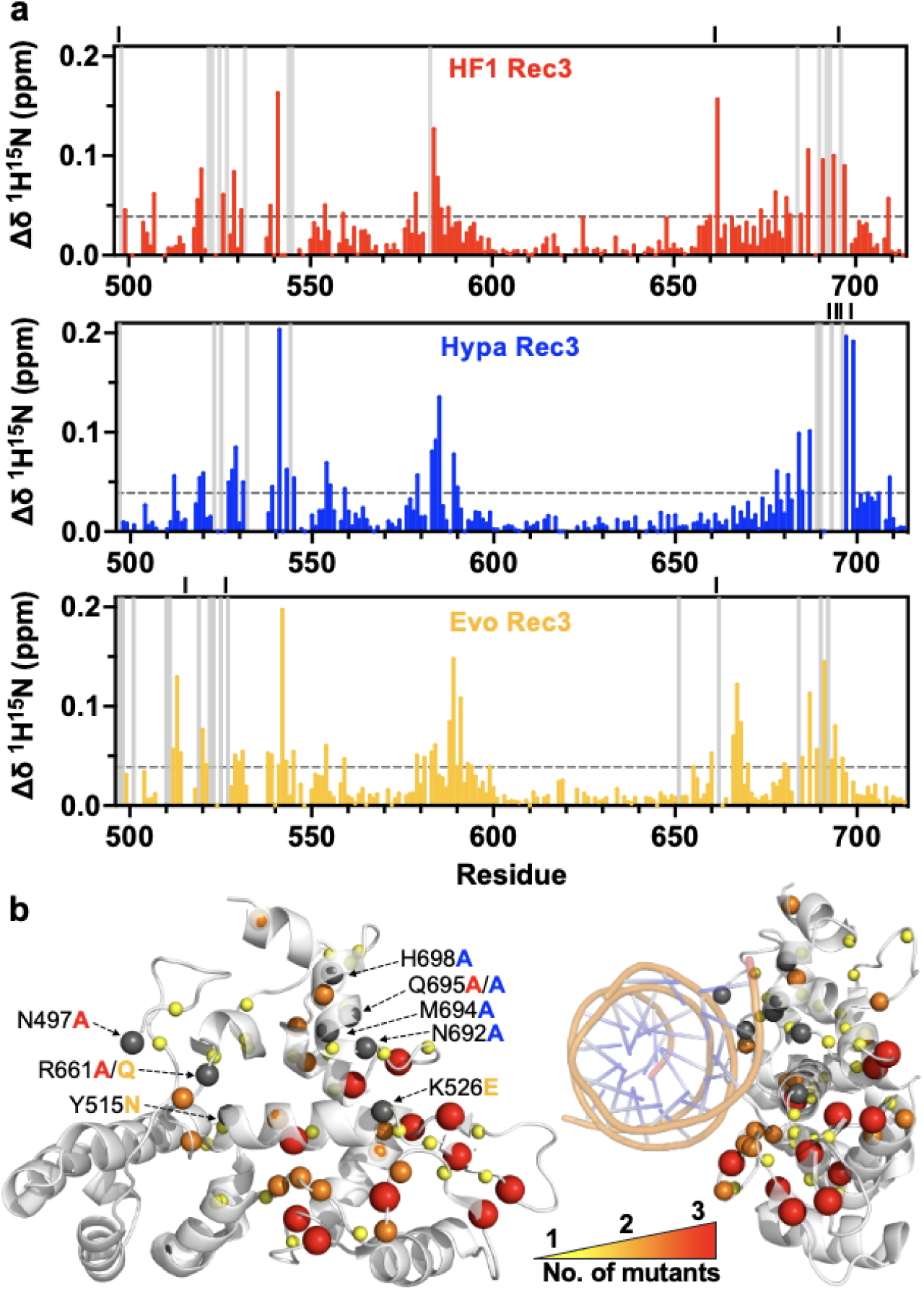
Chemical shift perturbations induced by high-specificity mutations reveal a structural signature in Rec3. **a.** ^1^H^15^N combined chemical shift perturbations (Δδ) for HF1 Rec3 (red), Hypa Rec3 (blue), and Evo Rec3 (yellow). Gray vertical bars represent line-broadened resonances and grey dashed lines denote 1.5σ above the 10% trimmed mean of all data sets. Sites of mutation are indicated by black lines above each plot. **b.** Significant ^1^H-^15^N chemical shift perturbations (+1.5σ) are mapped onto the Rec3 structure, with spheres denoting significant shifts observed in any one mutant (yellow), any two mutants (orange), and all three mutants (red). Sites of mutation are indicated by grey spheres and the mutated residues are labelled according to the colour scheme in (A). The structure on the right shows the RNA:DNA hybrid for positional reference to the observed perturbations.

Consistent patterns of chemical shift perturbation in all three variants face the RNA:DNA heteroduplex, and are localized around a loop region in proximity to the PAM-distal end of the hybrid and distant from the neighbouring domains. Such a structural change may alter points of contact between Rec3 and the RNA:DNA hybrid. Further, this region is in close proximity to the 692-699 α-helix (α37 from PDB: 4UN3)^7^ that was previously shown to insert within the RNA:DNA hybrid upon off-target DNA binding, suggesting a possible proofreading role in mismatch recognition^29^. Altered hybrid contacts and perturbation of α37 may be critical for enhancing the specificity of these variants.

For a more comprehensive analysis of the structural changes induced by specificity-enhancing mutations to Rec3, the carbon chemical shifts for HF1 Rec3, Hypa Rec3, and Evo Rec3 were also determined by NMR. Predictions of the secondary structure via Cα/Cβ chemical shifts show a general conservation of secondary structural features among the variants, which is further supported by the strong overlap of CD spectra for WT Rec3 and the variants (Supplementary Fig. 1a, b). ^1^H-^13^Cα/β/O spectral overlays of the variants with WT Rec3 reveal global spectral changes for each carbon position, indicating that the backbone and side chains are impacted by specificity-enhancing mutations (Supplementary Fig. 5). These structural perturbations are consistent with thermal unfolding experiments that show modest changes to the structural stability of the variants (Supplementary Fig. 1c). Analysis of ^13^Cα/β/_O_ chemical shift perturbations shows similar trends across the Rec3 variants, with the strongest effects occurring in the loop region proximal to the PAM-distal end of the RNA:DNA hybrid, consistent with previously discussed ^1^H-^15^N chemical shift perturbations (Fig. 3 and Supplementary Fig. 6). Collectively, these findings suggest that the high-specificity mutations induce a universal Rec3 structural change at the PAM-distal ends of the RNA:DNA hybrid, which may promote the specificity-enhancing properties of the Cas9 variants.

### High-specificity mutations perturb the Rec3 dynamic landscape

Analysis of the Rec3 dynamics in the three high-specificity variants reveals that the variants exhibit differential flexibility on the fast and slow timescales, with an observed shift toward slower timescale motions (Fig. 4a and Supplementary Figs. 7-11). In fact, decreased molecular motion on the ps – ns timescale via *S*^2^ is observed in all three variants, relative to WT Rec3 (Fig. 4b and Supplementary Fig. 10). CPMG relaxation dispersion experiments also reveal a change in quantitative μs – ms motions in the Rec3 variants (Fig. 4c, Supplementary Fig. 11, and Supplementary Tables 1-4). HF1 Rec3 exhibits fewer residues (−7 residues), Evo Rec3 exhibits slightly more residues (+4 residues), and Hypa Rec3 exhibits considerably more residues (+20 residues) with μs – ms flexibility than the WT Rec3. Interestingly, regardless of the number of residues exhibiting μs – ms motions, ∼50-60% of the sites identified in each variant are not found to undergo similar motions in WT Rec3, suggesting an overall rewiring of intradomain flexibility due to the specificity-enhancing mutations. In addition to a redistribution of slow timescale motions in Rec3, the residues that exhibit these motions in the variants have, on average, larger amplitudes to the CPMG profiles, reflecting a greater exchange contribution to relaxation (*R_ex_*) than observed in WT Rec3 (Fig. 4a and Supplementary Tables 1-4). HF1 Rec3 and Evo Rec3 display clusters of heightened flexibility at the RNA:DNA hybrid interface, while Hypa Rec3 enhances μs – ms motion across the entire domain, including regions interfacing with the RNA:DNA hybrid. These data reveal that high-specificity mutations perturb the Rec3 dynamic profile, particularly at the regions interfacing the RNA:DNA hybrid, which could remodel allosteric signalling within Cas9 through the interacting heteroduplex.

**Fig. 4.**
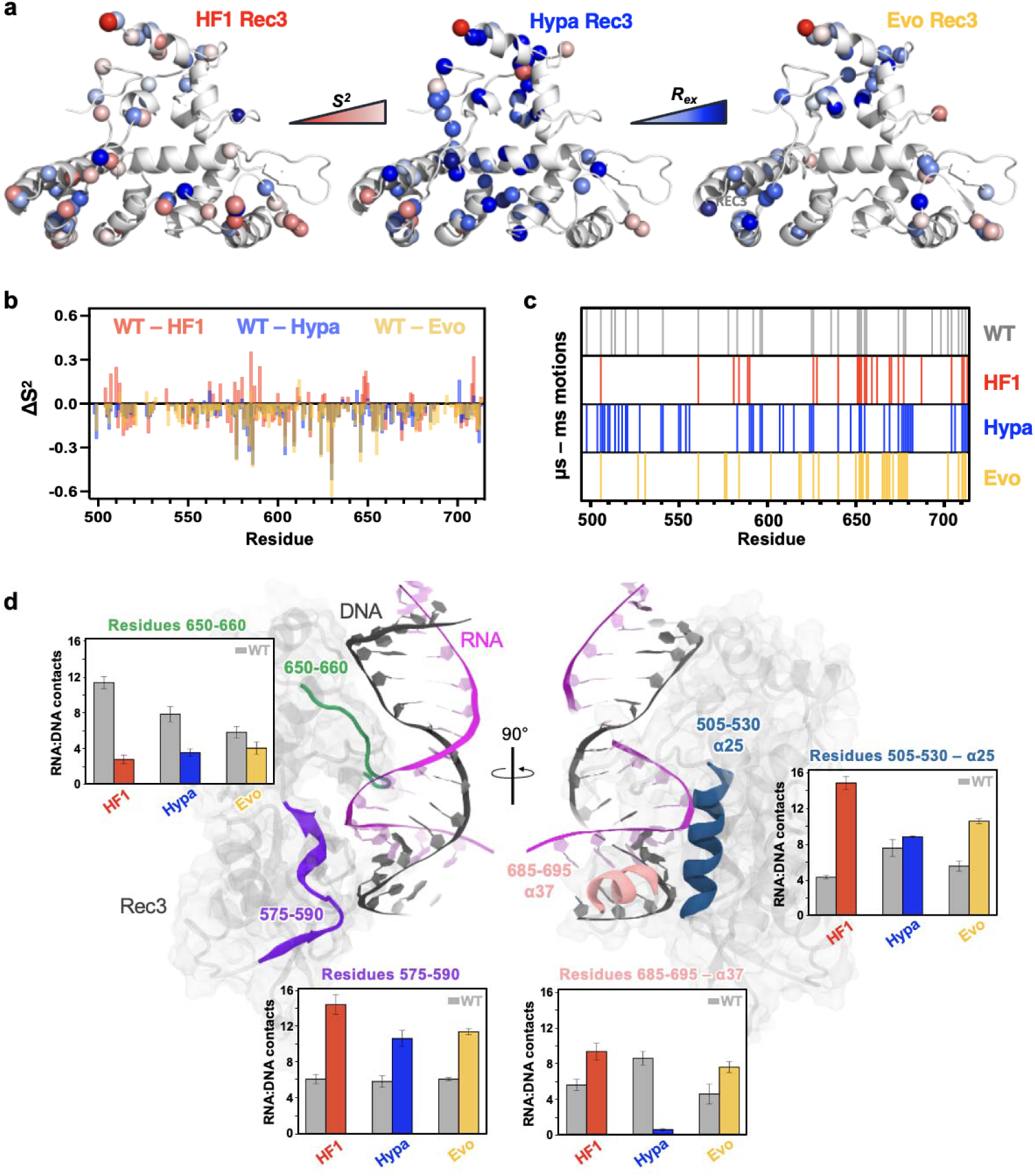
High-specificity mutations alter the dynamic profile of Rec3 and its interactions with the RNA:DNA hybrid. **a.** Residues exhibiting ps – ns motions via *S*^2^ and μs – ms motions via CPMG relaxation dispersion are mapped onto the Rec3 structure for HF1 Rec3 (left), Hypa Rec3 (centre), and Evo Rec3 (right). Fast motions were determined via *S*^2^ values 1.5σ below the 10% trimmed mean of all data sets. Red spheres represent ps – ns motions (*S*^2^), with a gradient denoting the lowest *S*^2^ values as dark red, signifying greater ps – ns flexibility. Blue spheres denote μs – ms motions (CPMG), with a blue gradient that becomes darker as the exchange contribution to relaxation (*R*_ex_) increases. **b.** Changes in *S*^2^ (Δ*S*^2^) between WT Rec3 and HF1 Rec3 (red), Hypa Rec3 (blue), and Evo Rec3 (yellow). Residues with positive Δ*S*^2^ values are more flexible on the ps – ns timescale in the variant, while residues with negative Δ*S*^2^ values are more flexible in WT. **c.** Residues exhibiting μs – ms motions via CPMG relaxation dispersion are indicated by a coloured bar for a per-residue comparison between the WT Rec3 (gray), HF1 Rec3 (red), Hypa Rec3 (blue), and Evo Rec3 (yellow). **d.** Differential contact analysis between the RNA:DNA hybrid and WT Rec3 (gray), HF1 Rec3 (red), Hypa Rec3 (blue), and Evo Rec3 (yellow). The number of stable contacts between Rec3 and the RNA:DNA hybrid is quantified in distinct regions of the protein, highlighted on the structure in blue (505-530), purple (575-590), green (650-660), and pink (685-695).

### High-specificity mutations in Rec3 alter interactions with the RNA:DNA hybrid

Given the structural perturbations observed experimentally, and the dynamic changes at the region comprising the interface of Rec3 and the RNA:DNA hybrid, we sought to assess whether the high-specificity mutations alter the interactions of the Rec3 domain with the hybrid through MD simulations. The difference in the Rec3–RNA:DNA hybrid contact stability between WT and the variants was estimated from the simulated ensemble (i.e., ∼16 μs for each system). A comparison of the total number of Rec3–hybrid contacts that are at least 10% more stable in either WT (grey) or the variants (red, blue, and yellow), shows noticeable mutation-induced changes (Fig. 4d). Specifically, the Rec3 regions comprising residues 512-530 (α-helix 25, α25 from PDB 4UN3^7^) and the 570-590 loop increase their contact stability with the RNA:DNA hybrid in all variants compared to the WT, with the α25 interactions remaining least perturbed by the HypaCas9 mutations. These regions also exhibit significant chemical shift perturbations (Fig. 3a). At the level of residues 692-699 (i.e., α37), Cas9-HF1 and evoCas9 display improved interactions with the hybrid with respect to WT Rec3, while a significant reduction is observed in HypaCas9, due to the non-polar alanine mutations exclusively present in this region (Fig. 4d). Notably, the α37 was shown to be involved in mismatch recognition, inserting within the RNA:DNA hybrid upon off-target DNA binding^29^. Resonances in the 692-699 region are also substantially line broadened or display CPMG relaxation dispersion, suggesting heightened flexibility (Figs. 3a, 4c). In light of this evidence, the different interactions of the α37 observed here for the three variants suggest a critical proofreading role.

Finally, at odds with the regions noted above, residues 650-660 show a consistent loss of stable contacts with the hybrid in the variants relative to the WT. Interestingly, this region does not exhibit NMR chemical shift perturbations (Fig. 3a). These results highlight important hotspots at the Rec3–hybrid interface, where rearranged molecular interactions with the key α37 might create the foundation for off-target detection.

### Mutations in Rec3 affect the HNH dynamics

Biochemical and single-molecule experiments indicated that Rec3 could act as an “allosteric effector” of HNH dynamics^3, 9, 13^. As the high flexibility of HNH ensures DNA cleavage at the proper location, alterations in the motions of HNH due to distal mutations in Rec3 could be critical for the specificity-conferring mechanisms of the variants studied here. We, therefore, sought to investigate whether the rewired intra-domain flexibility of Rec3 impacts the HNH domain. From the simulated ensembles of full-length Cas9 and its variants, we estimated differences in the HNH conformational dynamics by computing the Jensen-Shannon distance (JSD), which measures how similar two distributions are, ranging from 0 (similar distribution) to 1 (dissimilar distribution)^30^. The JSD was computed between the distributions of the intra-backbone distances (*JSD_BB-dist_*) and side-chain torsions (*JSD*_*sc-tor*_) of the HNH domain in the WT and variants (Fig. 5a). We observe that the HNH dynamics in evoCas9 are comparable to the WT for each estimated feature. On the other hand, in Cas9-HF1 and HypaCas9, the HNH dynamics are substantially reduced in terms of BB distances and SC torsions. Calculation of the differential root-mean-square fluctuations (ΔRMSF, Å) of the C⍰ atoms between the WT and variants (Fig. 5b) further shows that HNH flexibility is reduced by the HF1 and Hypa mutations, but remains comparable to WT in evoCas9.

**Fig. 5.**
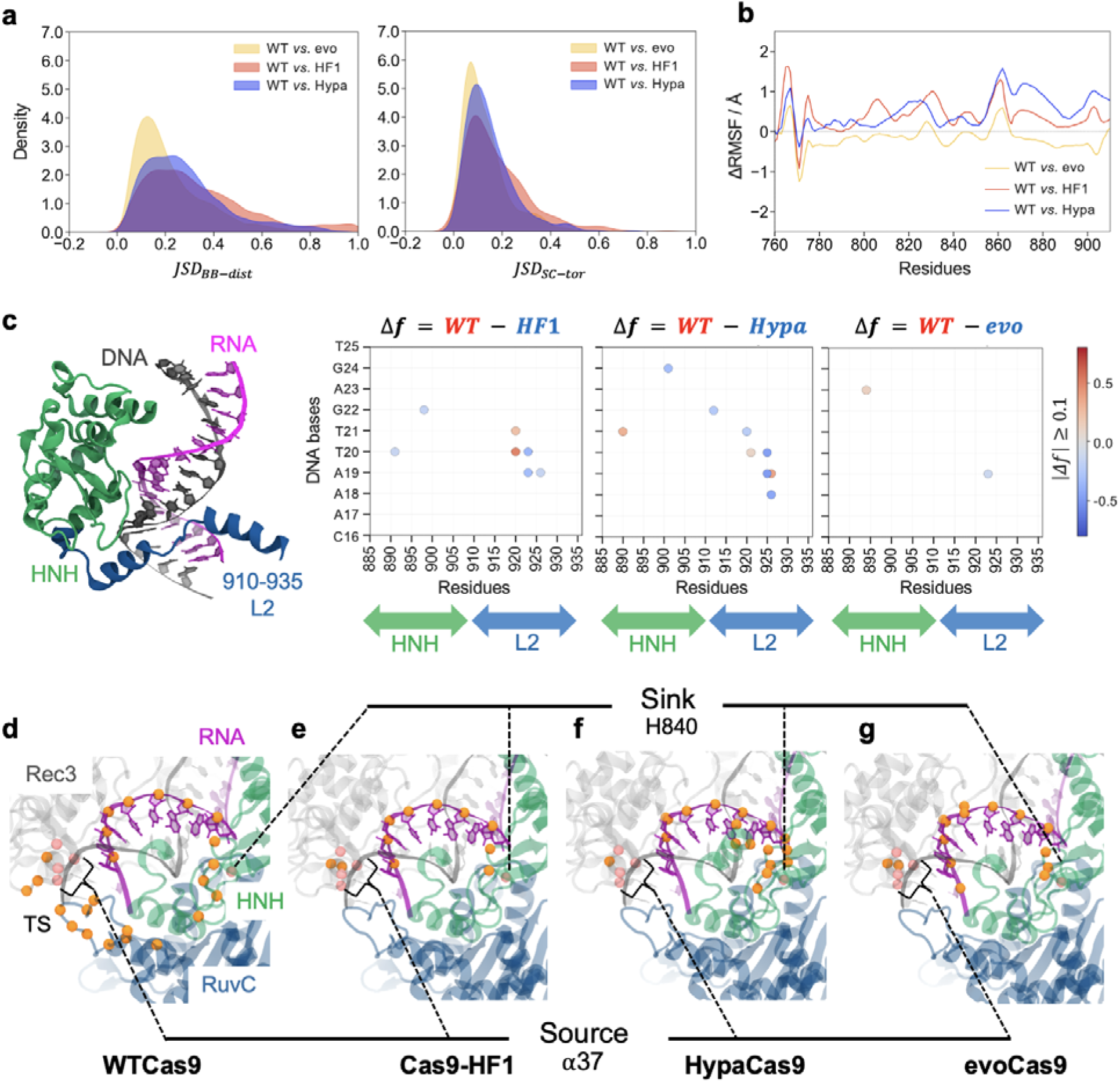
High-specificity mutations alter the HNH dynamics and reroute the allosteric signal in Cas9. **a.** Distribution of Jensen-Shannon Distances (JSD), comparing intra-backbone distances () and side-chain torsions () of the HNH domain in WTCas9 *vs.* Cas9-HF1 (red), WTCas9 *vs.* HypaCas9 (blue) and WTCas9 *vs.* evoCas9 (yellow). **b.** Differential root-mean-square fluctuations (ΔRMSF, Å) of the HNH ClZ atoms, computed between the WTCas9 and the HF1 (red), Hypa (blue), and Evo (yellow) variants. **c.** Interactions between the RNA:DNA hybrid and the HNH domain. Differential contact maps report residue pairs that gain contact stability in either the WTCas9 (red) or the HF1, Hypa, and Evo variants (blue). Residues 910-935 of HNH, which comprises the L2 loop, are indicated on the x-axis. The differential contact stability (Δ*f*) is computed as the difference in the frequency *f* of formed contacts in the WTCas9 and its variants and is plotted as |Δ*f*| ≥ 0.1, to account for contacts that are more stable in one system (e.g., WTCas9) by more than 10 % with respect to the other (e.g., variant). **d-g.** Signalling pathways connecting the α-helix 37 in Rec3 (i.e., source) to the HNH catalytic residue H840 (i.e., sink) in the WTCas9 (**d**), and the Cas9-HF1 (**e**), HypaCas9 (**f**), and evoCas9 (**g**) variants. Spheres of different colours are used to indicate the allosteric source/sink (pink) and the path nodes (orange). Analysis was performed on an aggregate sampling of ∼16 μs (i.e., four MD replicas of ∼4 μs each) for each system. Details are in the *Materials and Methods* section and in Supplementary Fig. 13.

Principal Component Analysis (PCA) also shows that Evo HNH samples a conformational space more similar to the WT HNH, than to HF1 and Hypa HNH (Supplementary Fig. 12). We next analysed the interactions between the heteroduplex and the HNH domain in the three variants, thereby gaining insight into whether the RNA:DNA hybrid, interposing between Rec3 and HNH, could transduce the flexibility of Rec3 to the HNH domain. The gain and loss of pairwise interactions between the RNA:DNA hybrid bases and the HNH residues were estimated as the number of contacts between pairs that are at least 10% more stable in the WT or the variants (details in the *Supplementary Materials and Methods*). Interestingly, interactions between the DNA target strand bases and the residues 910-935 of HNH, which comprise the L2 loop, are increased in Cas9-HF1 (Fig. 5c). HypaCas9 also displays an increase in DNA-L2 interactions compared to the WT, while these interactions are least perturbed in evoCas9. The L2 loop of HNH was previously shown to exert a critical role in mismatch recognition in WTCas9, as it tightly binds mismatched DNA bases^29^. This reduces the conformational mobility of HNH required for its transition to the active state, thus exerting a regulatory mechanism. In Cas9-HF1 and HypaCas9, the increase in interactions between the DNA and L2 is associated with altered dynamics of HNH (from the JSD plots). On the other hand, in evoCas9, where these interactions are less affected, the HNH dynamics are comparable to those of WTCas9. This observation suggests that the RNA:DNA hybrid transduces the effect of the Rec3 mutations to HNH through its interaction with the L2 loop.

### Rewired allosteric pathways between Rec3 and HNH

The perturbations in the Rec3 domain and associated changes to HNH dynamics observed via MD simulations hint at rewired allosteric pathways between the two domains, induced by the mutations. To gain insight into this communication, we performed shortest path calculations between the Rec3 mutation sites and the HNH catalytic core based on a dynamic network model from the simulated ensemble (see Materials and Methods). For each pair of “source” (Rec3 mutation sites) and “sink” (HNH catalytic residue H840), the optimal path (i.e., the shortest) as well as the top five sub-optimal paths were computed. Then, the distribution of all path lengths (i.e., the number of edges connecting the source and sink) corresponding to the optimal and sub-optimal paths, were estimated, alongside the occurrence of residues in the paths to check for convergence in the communication pathways (Supplementary Figs. 13-15).

We observe an interesting trend in the communication pathways sourcing from residues located within the critical α37. In the WTCas9, α37 communicates with the HNH catalytic core via two main routes passing through the guide RNA and the RuvC residues (Fig. 5d, Supplementary Fig. 13). In the Cas9-HF1, paths sourcing from α37 pass almost exclusively through the guide RNA (Fig. 5e), thereby favouring the signal transduction. In this system, the communication routes that source from residue 695, which is within α37, also display a significant reduction in the path lengths with respect to the WT, while paths involving the residues 497 and 661 are less affected (Supplementary Fig. 14a). In HypaCas9, where all Rec3 mutations are clustered in α37 (N692A, M694A, Q695A, H698A), the RuvC-mediated path observed in the WT is abolished, increasing the occurrence of the RNA nucleotides in the Rec3-HNH communication (Fig. 5f). Path lengths from this region to the HNH core are reduced relative to the WT, and communication routes also converge noticeably (Supplementary Fig. 13). We interestingly observe that common mutation sites at the level of α37 display similar communication patterns in the HF1 and Hypa variants, resulting in the replacement of the RuvC-meditated path (observed in the WT) with guide RNA nucleotides. We therefore sought to analyse the routes connecting α37 to the HNH core in the evoCas9 variant, which does not hold mutations in this α-helix. Interestingly, the Evo mutations also abolish the RuvC-mediated pathway, re-routing the communication to the guide RNA, and altering the WT Rec3-HNH communication similar to the HF1 and Hypa mutations located within α37 (Fig. 5g). On the other hand, pathways sourcing from the Evo mutation sites are preserved when moving from the WT to the Evo variant, and path lengths are not significantly affected (Supplementary Fig. 15).

The three variants thereby alter the communication between α37 in Rec3, implicated in DNA mismatch recognition^29^, and the HNH catalytic core. While HypaCas9 exerts this effect directly through mutations located within the key α-helix, Cas9-HF1 uses the local Q695A mutation, alongside distal mutations, to re-route the signal. evoCas9 most strikingly leverages purely distal mutations to allosterically affect the communication paths between the α37 and the HNH core. In this respect, it is notable that Cas9-HF1 and evoCas9 share mutations in residue 661, distal to the key α-helix. An in-depth analysis of the paths sourcing from the α37 also shows that the path lengths are reduced in all variants with respect to the WT (Supplementary Fig. 13), suggesting that the three variants enhance communication from the DNA mismatch recognition α-helix 37 to the HNH active site, transferring the information of mismatch tolerance to the active site.

## DISCUSSION

We combined solution NMR with molecular simulations to decipher the dynamics and allostery of three high-specificity variants of the Cas9 enzyme – Cas9-HF1, HypaCas9, and evoCas9 – that contain multiple mutations within the Rec3 domain. Solution NMR was used to measure the structure and dynamics of the Rec3 domain with experimental accuracy, and MD simulations provided an all-atom description of the allosteric phenomenon, and how it propagates from Rec3 within the full-length complex. We detect conserved structural features and dynamic differences in the HF1, Hypa, and Evo variants, which rewire the Rec3 domain flexibility and remodel the allosteric signalling of Cas9. We found a consistent structural change in the Rec3 domain induced by the high-specificity mutations (Fig. 3a), with NMR chemical shift perturbations observed on the face of Rec3 that interfaces with the RNA:DNA hybrid at the PAM distal ends. Solution NMR also revealed altered molecular motions that perturb the Rec3 dynamic profile, particularly at regions interfacing with the heteroduplex. Relaxation dispersion experiments revealed μs – ms dynamics across the Hypa Rec3 domain, including at the RNA:DNA hybrid interface (Fig. 4a,c), while HF1 and Evo Rec3 also show clusters of heightened flexibility near the interface. These structural and dynamic changes suggest that high-specificity mutations could affect the interactions of the Rec3 domain with the heteroduplex. This is also supported by single-molecule data, showing that the Rec3 mutations interfere with the RNA:DNA heteroduplexation process,^17^ and with DNA unwinding/rewinding^15, 16^. The Rec3–hybrid contacts were thereby obtained from multi-μs MD simulations of the full-length CRISPR-Cas9 systems (Fig. 4d). The three variants exhibited similar trends in altered interactions with the RNA:DNA hybrid at the level of the 575-590 and 650-660 loops, and the α25 helix. Most notably, they differentially interact with the heteroduplex at the level of α37 (Fig. 4b), which was shown to “sense” DNA mismatches within the RNA:DNA hybrid and is critically involved in mismatch recognition^29^. In HypaCas9, which holds all of its mutations within α37, interactions with the hybrid are diminished relative to the WTCas9. Conversely, increased interactions between α37 and the hybrid are detected in the HF1 and Evo variants.

Critically, the high-specificity mutations in Rec3 also allosterically affect the dynamics of the catalytic HNH domain. Analysis of the interactions between HNH and the RNA:DNA heteroduplex reveals important differences at the level of the L2 loop (Fig. 5b), which was shown to tightly bind mismatched DNA bases in WTCas9, reducing the conformational mobility of HNH as a regulatory mechanism^29^. In HypaCas9, where the key α37 in Rec3 detaches from the hybrid, L2 in HNH tightly binds the hybrid. In Cas9-HF1 and evoCas9, where α37 has stronger interactions with the hybrid, the DNA-L2 interactions are less affected than observed for HypaCas9, though still increased relative to WTCas9. Consistent with the increased L2-hybrid interactions, the three variants affect the dynamics of the catalytic HNH domain (Fig. 5a-b), which exhibits reduced flexibility from the WT in Cas9-HF1 and HypaCas9, while remaining comparable in evoCas9. This aligns with single-molecule studies of HF1 and Hypa, showing that distal mutations in Rec3 alter the conformational equilibrium of HNH^3^, and is interesting in light of biochemical studies profiling the trade-off between activity and specificity in the Cas9 variants^31^. Though the variants exhibit comparable specificity for off-target detection, Cas9-HF1 and HypaCas9 have significantly higher on-target activity than evoCas9. As Cas9 activity can be a direct consequence of HNH dynamics^32^, the alterations to the motions of HNH, observed in the HF1 and Hypa variants in this study and through single-molecule experiments^3^, could be a crucial component to maintaining their on-target activity while also conferring high specificity.

We also note that solution NMR detected conserved structural features in the HF1, Hypa, and Evo Rec3 domains, irrespective of the locations of their high-specificity mutations (Fig. 3). These structural changes, mainly found in regions of Rec3 interfacing with the RNA:DNA hybrid, could constitute a signature for the comparable specificity of the three variants^31^. Alongside these conserved structural changes, dynamical differences in the three variants involve two critical elements of mismatch tolerance, – α37 in Rec3 and L2 in HNH – that impact important contacts with the RNA:DNA hybrid and modulate Rec3 and HNH dynamics (Figs. 4-5). Intriguingly, shortest-path analysis reveals that the variants rewire communication between α37 of Rec3 and the HNH catalytic core in a consistent manner (Fig. 5c). In fact, the variants reroute the communication through the guide RNA, reducing the path lengths and increasing the communication efficiency with respect to WTCas9. HypaCas9, which holds its mutations within α37, directly alters the signal transfer, while Cas9-HF1 uses the α37 Q695A mutation and its other distal mutations, and evoCas9 fully leverages mutations that are not located within α37, to reroute the signal from this critical α-helix to the HNH core via protein dynamics. These data are consistent with our experimental findings, where the disparate mutations of the variants all induce structural perturbations and dynamic line broadening at α37 (Fig. 3), either directly or allosterically, while shifting the flexibility of the domain toward slower molecular motions distinct from WT Rec3. These data suggest an allosteric mechanism in which the three high-specificity Cas9 variants increase communication between the DNA mismatch recognition helix and the HNH active site. This outcome relies on subtle structural and dynamic perturbations to the WT Rec3 domain that transfers the information of mismatch tolerance to the active site.

## CONCLUSIONS

In summary, solution NMR and MD simulations reveal that the Cas9-HF1, HypaCas9, and evoCas9 high-specificity mutations alter the structure and dynamics of the Rec3 domain, and remodel the allosteric signalling within Cas9. High-specificity mutations induce conserved structural changes at regions of Rec3 that interface with the RNA:DNA hybrid, and subtle dynamic variations that remodel the communication from the DNA recognition region to the HNH catalytic core. In the three variants, the Rec3 domain senses the RNA:DNA hybrid, whose interaction transduces the allosteric signal from Rec3 to HNH. These interactions are associated with altered dynamics of the HNH domain, which is more pronounced in Cas9-HF1 and HypaCas9, in agreement with single-molecule data. The three variants consistently reroute the communication between the α-helix 37 in Rec3, previously shown to sense target DNA complementarity and the HNH catalytic core. While HypaCas9 exerts this effect directly through mutations within α37, Cas9-HF1 uses the local Q695A mutation, alongside its other mutations. Most strikingly, evoCas9 fully leverages distal mutations to allosterically reroute the signal. These findings highlight a mechanism in which the three disparate Cas9 variants increase the communication from the DNA mismatch recognition helix to the HNH core through structural and motional perturbations, transferring the information of mismatch tolerance to the nuclease active site.

## MATERIALS AND METHODS

### Protein Expression and Purification

The WT Rec3 domain (residues 497-713) of the *S. pyogenes* Cas9 was engineered into a pET-28a(+) vector with an N-terminal His_6_-tag and a TEV protease cleavage site. High-specificity Rec3 variants, including HF1 Rec3 (N497A, R661A, Q695A), Hypa Rec3 (N692A, M694A, Q695A), and Evo Rec3 (Y515N, K526E, R661Q), were generated via mutagenesis. The WT and high-specificity Rec3 plasmids were transformed into BL21-Gold (DE3) competent cells (Agilent) and expressed in lysogeny broth (LB) for X-ray crystallography and biochemical studies, and deuterated M9 minimal media, isotopically enriched with ^15^NH_4_Cl and ^13^C_6_H_12_O_6_ (Cambridge Isotope Laboratories) as the sole nitrogen and carbon sources and supplemented with MgSO_4_, CaCl_2_, and MEM vitamins, for NMR experiments. Cells were grown at 37°C to an OD_600_ of 0.6 – 0.8 (LB) or 0.8 – 1.0 (deuterated M9), induced with 1.0 mM IPTG, incubated for 16-18 hours at 20 °C, and then harvested by centrifugation.

During purification of WT Rec3 and the Rec3 variants, cells were resuspended in a buffer of 20 mM sodium phosphate, 300 mM sodium chloride, and 5 mM imidazole at pH 8.0 with a mini EDTA-free protease inhibitor cocktail tablet (Sigma Aldrich) and lysed by sonication. Following removal of cell debris by centrifugation, Rec3 was purified from the supernatant by Ni-NTA affinity chromatography. The column was washed with lysis buffer and the protein was eluted with a buffer of 20 mM sodium phosphate, 300 mM sodium chloride, and 250 mM imidazole at pH 8.0. The N-terminal His-tag was cleaved by TEV protease for 4 hours at room temperature while being dialyzed into lysis buffer and subsequently removed by Ni-NTA chromatography. Rec3 was further purified by size exclusion chromatography via a HiLoad 26/600 Superdex 200 column (GE) in a buffer of 20 mM sodium phosphate, 300 mM sodium chloride, and 2 mM EDTA at pH 8.0. Fractions containing purified Rec3 were then dialyzed into a buffer containing 40 mM sodium phosphate, 50 mM potassium chloride, 1 mM EDTA, and 6% D_2_O at pH 7.5, and concentrated to 0.65 mM for NMR experiments.

### Nuclear Magnetic Resonance (NMR) Spectroscopy

NMR experiments were performed on Bruker Avance NEO 600 MHz and Bruker Avance III HD 850 MHz spectrometers at 25°C. NMR data were processed using NMRPipe^33^ and analysed in Sparky^34^ with the help of in-house scripts. Backbone resonances for WT Rec3 were previously assigned (BMRB entry 50389)^35^ and transferred to the HSQC spectra of the Rec3 variants. ^1^H^15^N combined chemical shift perturbations were determined from ^1^H^15^N TROSY HSQC^36^ spectra by 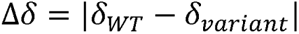, where 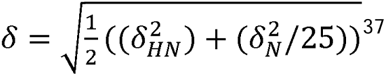. Carbon chemical shifts (^13^Cα/β/O) were obtained from HNCA, HNCACB, and HNCO experiments on each variant, and ^13^C chemical shift perturbations were determined by Δ*c = |δ_WT_* -*δ_variant_*|, where δ are the ^13^C chemical shift values in ppm.

TROSY-based spin relaxation experiments were performed with the ^1^H and ^15^N carriers set to the water resonance and 120 pm, respectively. Longitudinal relaxation rates (*R_1_*) were measured with *T*_1_ delays of 0, 20(x2), 60(x2), 100, 200, 600(x2), 800, and 1200 ms. Transverse relaxation rates (*R_2_*) were measured with *T_2_* delays of 0, 16.9, 33.9 (x2), 67.8, 136 (x2) and 203(x2) ms. The recycle delay in these experiments was 2.5 s^38^ and these data were collected in a temperature-compensated manner. Longitudinal and transverse relaxation rates were extracted by nonlinear least squares fitting of the peak heights to a single exponential decay using in-house software. Uncertainties in these rates were determined from replicate spectra with duplicate relaxation delays. Steady-state ^1^H-[^15^N] NOE were obtained by interleaving pulse sequences with and without proton saturation and calculated from the ratio of peak heights^38^. For calculation of order parameters, model-free analysis was carried out using the Lipari-Szabo formalism in RELAX^39^ with fully automated protocols^40^. Carr-Purcell-Meiboom-Gill (CPMG) NMR experiments were adapted from the report of Palmer and coworkers^41^, and performed at 25°C with a constant relaxation period of 40 ms, a 2.0 second recycle delay, and ν_CPMG_ values of 25, 50 (x2), 75, 100, 150, 250, 500 (x2), 750, 800 (x2), 900, 1000 Hz. Relaxation dispersion profiles were generated by plotting *R*_2_ vs. 1/τ_cp_ and exchange parameters were obtained from fits of these data using NESSY^42^. Profiles were fit to three models, with the best model selected using Akaike information criteria with second order correction.

Model 1: No exchange

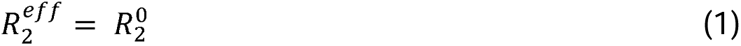

Model 2: Two-state, fast exchange (Meiboom equation^43^)

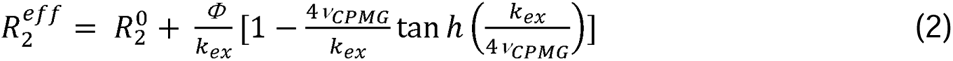

Model 3: Two state, slow exchange (Richard-Carver equation^44^)

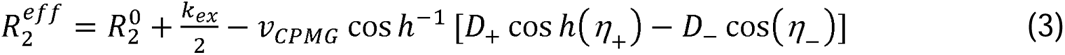

### Circular Dichroism (CD) Spectroscopy

Rec3 samples (10 μM) were dialyzed into a buffer containing 40 mM sodium phosphate, 1 mM EDTA at pH 7.5 for CD experiments. CD spectra and thermal unfolding experiments were collected on a JASCO J-815 spectropolarimeter equipped with a variable temperature Peltier device and using a 2 mm quartz cuvette. CD spectra were collected at 25°C, and denaturation curves were recorded at 208 nm over a temperature range of 20 – 80°C. *T_m_* values were determined via nonlinear curve fitting in GraphPad Prism.

### X-ray Crystallography

WT Rec3 was crystallized at room temperature by sitting drop vapor diffusion. Rec3 protein at 12 mg/mL in 20 mM Tris pH 7.5, 50 mM potassium chloride, 1mM EDTA was mixed with crystallizing condition 0.1 M Bis-Tris pH 5.5, 0.2 M sodium chloride, 25% PEG3350 at a 2:1 ratio of protein-to-crystallizing condition. Crystals were cryoprotected in 30% ethylene glycol diluted in crystallizing condition. Diffraction images were collected at the NSLS-II AMX beamline at Brookhaven National Laboratory. Images were processed using XDS^45^ and Aimless^46^ in CCP4 and the structure was solved by molecular replacement with Phaser in Phenix^47^. The region of the full-length SpCas9 crystal structure corresponding to the Rec3 domain (residues 501-710, PDB ID: 4UN3)^7^ was used as the search model for molecular replacement. The Rec3 structure was finalized by iterative rounds of manual building in Coot^48^ and refinement with Phenix.

### Molecular Dynamics (MD) Simulations

MD simulations were performed on the full-length CRISPR-Cas9 systems, based on the X-ray structure of the *S. pyogenes* Cas9 (PDB: 4UN3, at 2.59 Å resolution)^7^. This structure captures the HNH domain in an inactivated state, a so-called “conformational checkpoint” between DNA binding and cleavage^9^, in which the RNA:DNA complementarity is recognized before the HNH domain assumes an activated conformation. Single-molecule experiments reported alterations of the HNH dynamics in this “conformational checkpoint” in the high-specificity Cas9 variants^3^, which motivated the use of this structure for MD simulations in the present study. Four model systems were considered: the WTCas9, Cas9-HF1, HypaCas9, and evoCas9. Each system was solvated in a ∼145 × ∼110 × ∼147 Å periodic box of ∼220,000 total atoms. In all systems, counterions were added to provide physiological ionic strength.

The systems were simulated using the AMBER ff99SBnmr2 force field for the protein^49^, which improves the consistency of the backbone conformational ensemble with NMR experiments, also used in our previous NMR/MD study^50^. Nucleic acids were described using the ff99bsc1 corrections for DNA^51^ and the *χ*OL3 corrections for RNA^52, 53^. The TIP3P model was used for water^54^. An integration time step of 2 fs was used. All bond lengths involving hydrogen atoms were constrained using the SHAKE algorithm. Temperature control (300 K) was performed via Langevin dynamics^55^, with a collision frequency γ = 1/ps. Pressure control was achieved through a Berendsen barostat^56^, at a reference pressure of 1 atm and with a relaxation time of 2 ps. The systems were subjected to energy minimization to relax water molecules and counterions, keeping the protein, RNA, and DNA fixed with harmonic positional restraints of 300 kcal/mol · Å^2^. The systems were heated from 0 to 100 K in the canonical NVT ensemble, by running two simulations of 5 ps each, imposing positional restraints of 100 kcal/mol · Å^2^ on the above-mentioned elements of the systems. The temperature was further increased to 200 K in ∼100 ps of MD in the isothermal-isobaric NPT ensemble, reducing the restraint to 25 kcal/mol Å^2^. Then, all restraints were released and the temperature was raised to 300 K in a single NPT run of 500 ps. Finally, ∼10 ns of NPT runs allowed the density of the system to stabilize around 1.01 g/cm^-3^. Production runs were carried out in the NVT ensemble reaching ∼4 µs for each system, and in four replicates, collecting ∼16 µs per system (for a total of ∼64 μs of simulation time). This multi-µs simulation length with replicates was motivated by our previous work^10, 50, 57^, showing that it provides a solid statistical ensemble for the analysis of allosteric mechanisms (described below). All simulations were conducted using the GPU-empowered version of AMBER 20^58^. Analysis of the results was performed on the aggregate ensemble (i.e., ∼16 μs per system), after discarding the first ∼200 ns of MD from each trajectory, to enable proper equilibration and fair comparison.

### Analysis of Jensen-Shannon Distances

To characterize the difference in the conformational dynamics of the HNH domain, we analysed the distributions of all intra-backbone distances (*BB* – *dist*) side-chain torsions (*SC* – *tor*) in the investigated systems. To compare the distributions of the abovementioned features between any two of our systems, we computed the Jensen-Shannon Distance (*JSD* or *D_JS_*), a symmetrized version of Kullback-Leibler divergence (*D_KL_*)^30^. The *JSD* ranges from 0 to 1, where 0 corresponds to two identical distributions and 1 corresponds to a pair of separated distributions. For two distributions *P_i_* and *P_j_*, and considering a feature *x_f_* from two different ensembles *i* and *j*,

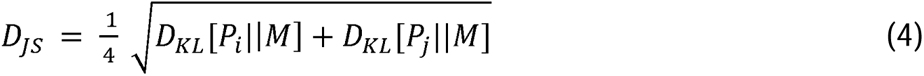

where 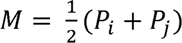. The Kullback-Leibler divergence, *D_KL_*, corresponds to two distributions *P_i_* and *P_j_* is of the following form:

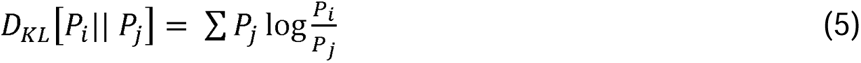

*JSD* values were computed using the Python ENSemble Analysis (PENSA) open-source library ^59^. Kernel density estimations of the *JSD* values were plotted to describe the *JSD* range and compare the systems.

### Shortest-paths calculation

The allosteric pathways for information transfer were characterized through dynamic network analysis^60^, a well-established method for the study of allosteric effects in proteins and nucleic acids^61–63^. Through this analysis, the CRISPR-Cas9 system, and its HF1, Hypa, and Evo variants, are described as graphs of nodes and edges, where nodes represent the amino acid (Cα atoms) and the nucleotides (P atoms, N1 in purines and N9 in pyrimidines)^64^, and edges denote the connection between them. To account for the information exchange between amino acids and nucleobases, the length of the edges connecting nodes was related to their motion correlations. A Generalized Correlations (GC) method^65^ was used, which quantifies the system’s correlations based on Mutual Information, describing the correlation between residue pairs independently on their relative orientation and capturing non-linear contributions to correlations (details on GC analysis are reported in the *Supplementary Materials and Methods*).

Based on GC analysis, the weight (*w_ij_*) of the edges connecting nodes *i* and *j* was computed as:

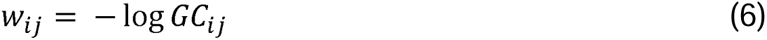

placing strongly correlated nodes close to each other in the dynamical network (i.e., displaying shorter edge lengths). To determine which nodes are in “effective contact”, two nodes were considered connected if any heavy atom of the two residues is within 5 A□ of each other (i.e., distance cut-off) for at least 70 % of the simulation time (i.e., frame cut-off). This threshold was shown to properly optimize the network structure in our previous studies of CRISPR- Cas9^10, 50, 57^. The resulting “weighted graph” defines a dynamical network, used for shortest-paths calculation.

Shortest-paths calculations were carried out computing the optimal (i.e., the shortest) and top five sub-optimal pathways between distally separated sites. Indeed, although the optimal path corresponds to the most likely mode of communication between nodes, suboptimal paths can also be crucial for the transmission of allosteric communication by providing alternative routes^60^. Hence, in addition to the optimal path, we also considered the top five sub-optimal pathways.

Well-established algorithms were employed for shortest-paths analysis. The Floyd-Warshall algorithm^66^ was utilized to compute the optimal paths between the network nodes. This algorithm first creates a matrix of dimension *n* x *n*, where *n* is the number of nodes. The elements of the matrix are the distance between all vertices in the network. The algorithm then recursively checks, for each pair of nodes (e.g., *i* and *j*), if there’s any intermediate node (e.g., *k*) such that the sum of distances between *i* - *k* and *k* - *j* are less than the current distance of *i* - *j*. For each pair of nodes, the algorithm reiterates its checks as many times as the number of nodes in the graph. The resulting final matrix contains the shortest distance between all the vertices in the graph. The five sub-optimal paths were computed in rank from the shortest to the longest, using Yen’s algorithm^67^, which computes single-source *K*-shortest loop-less paths (i.e., without repeated nodes) for a graph with non-negative edge weights.

For each system, the shortest pathways were computed between each mutation site in Rec3 (defined as source nodes) and the HNH catalytic residue H840 (defined as sink node). In Cas9-HF1, the shortest pathways were computed sourcing from the N497A, R661A, and Q695A. In evoCas9, shortest-paths calculations were sourced from the M495V, Y515N, K526E, and R661Q mutations. In HypaCas9, where all Rec3 mutations are clustered in the α-helix 37 (N692A, M694A, Q695A, H698A), the distribution of the path lengths was plotted by combining all the pathways connecting the N692A, M694A, Q695A, and H698A mutation sites to the H840 catalytic site. For comparison, shortest-paths calculations were also performed in the WTCas9, defining the same source and sink nodes of the variants. Given the importance of the α-helix 37 for mismatch recognition^29^, we also performed shortest-paths calculations sourcing from the HypaCas9 mutations, which locate within this key α-helix, in the Cas9-HF1 and evoCas9 variants, as well as in the WTCas9. To compare the systems, the distribution of the path lengths was plotted by combining all the pathways connecting all HypaCas9 mutation sites to H840.

All shortest-paths calculations were performed over ∼16 μs of aggregate sampling for each system (arising from four simulation replicates). To compare the lengths of the paths among the systems, we plotted the kernel density estimates (KDE) as a function of the path lengths (i.e., the number of edges connecting source and sink) for each of the source-sink pairs. To check for the convergence of communications in the systems, we estimated the occurrence of residues falling in the optimal and top five sub-optimal pathways. For each system, the aggregate ensemble arising from four simulation replicates (i.e., ∼16μs) was sectioned into three sample pools, which were used to compute the average occurrences and estimate the associated standard error of the mean. Networks of all the systems were built using the Dynetan Python library^60^. Path-based analyses were performed using the NetworkX Python library^68^ and our in-house Python scripts (available at: https://github.com/palermolab).

## Acknowledgments

This material is based upon work supported by the National Institutes of Health (Grant No. R01GM136815 to G.P.L. and G.P.; and R01GM141329, to G.P.) and the National Science Foundation (Grant No. MCB 2143760 to G.P.L. and CHE-2144823, and CHE-1905374 to G.P.). Part of this work used Expanse at the San Diego Supercomputer Center through allocation MCB160059 and Bridges2 at the Pittsburgh Supercomputer Center through allocation BIO230007 from the Advanced Cyberinfrastructure Coordination Ecosystem: Services & Support (ACCESS) program, which is supported by National Science Foundation grants #2138259, #2138286, #2138307, #2137603, and #2138296. Computer time was also provided by the National Energy Research Scientific Computing Center (NERSC) under Grant No. M3807. This work also used the AMX beamline of the National Synchrotron Light Source II, a U.S. Department of Energy (DOE) Office of Science user facility operated for the DOE Office of Science by Brookhaven National Laboratory under Contract No. DE-SC0012704. The Center for BioMolecular Structure (CBMS) is primarily supported by the National Institute of General Medical Sciences through a Center Core Grant (No. P30GM133893), and by the DOE Office of Biological and Environmental Research (Grant No. KP1605010).

## Author Contributions

ES expressed and purified Cas9 proteins, performed NMR and CD experiments, and analyzed the data; SS and MA performed MD simulations and analyzed the data; AMD performed X-ray crystallography and analyzed the data; GJ supervised X-ray crystallography; GP conceived the project and supervised MD simulations; GPL conceived the project and supervised NMR studies; ES, SS, GP, and GPL wrote the initial draft and the final manuscript was written with contributions from all authors.

## Data Availability Statement

Data are available from the corresponding authors upon reasonable request.

## Additional Information

Additional information is available as supplementary information.

## REFERENCES

1. Doudna, J. A. The promise and challenge of therapeutic genome editing. Nature 578, 229–236 (2020).

2. Fu, Y. et al. High-frequency off-target mutagenesis induced by CRISPR-Cas nucleases in human cells. Nat. Biotechnol. 31, 822–826 (2013).

3. Chen, J. S. et al. Enhanced proofreading governs CRISPR–Cas9 targeting accuracy. Nature 550, 407–410 (2017).

4. Kleinstiver, B. P. et al. High-fidelity CRISPR–Cas9 nucleases with no detectable genome-wide off-target effects. Nature 529, 490–495 (2016).

5. Casini, A. et al. A highly specific spCas9 variant is identified by in vivo screening in yeast. Nat Biotechnol 36, 265–271 (2018).

6. Jiang, F. & Doudna, J. A. The structural biology of CRISPR-Cas systems. Curr. Opin. Struct. Bio.l 30, 100–111 (2015).

7. Anders, C., Niewoehner, O., Duerst, A. & Jinek, M. Structural basis of PAM-dependent target DNA recognition by the Cas9 endonuclease. Nature 513, 569–573 (2014).

8. Sternberg, S. H., Redding, S., Jinek, M., Greene, E. C. & Doudna, J. A. DNA interrogation by the CRISPR RNA-guided endonuclease Cas9. Nature 507, 62–67 (2014).

9. Dagdas, Y. S., Chen, J. S., Sternberg, S. H. & Doudna, J. A. A Conformational Checkpoint between DNA Binding and Cleavage by CRISPR-Cas9. Sci. Adv. 3, eaao002 (2017).

10. East, K. W. et al. Allosteric Motions of the CRISPR-Cas9 HNH Nuclease Probed by NMR and Molecular Dynamics. J. Am. Chem. Soc. 142, 1348–1358 (2020).

11. Jiang, F. et al. Structures of a CRISPR-Cas9 R-loop complex primed for DNA cleavage. Science 351, 867–871 (2016).

12. Palermo, G., Miao, Y., Walker, R. C., Jinek, M. & McCammon, J. A. Striking Plasticity of CRISPR-Cas9 and Key Role of Non-target DNA, as Revealed by Molecular Simulations. ACS Cent. Sci. 2, 756–763 (2016).

13. Yang, M. et al. The Conformational Dynamics of Cas9 Governing DNA Cleavage Are Revealed by Single-Molecule FRET. Cell Rep. 22, 372–382 (2018).

14. De Paula, V. S., Dubey, A., Arthanari, H. & Sgourakis, N. G. A slow-exchange conformational switch regulates off-target cleavage by high-fidelity Cas9. Preprint at https://www.biorxiv.org/content/10.1101/2020.12.06.413757 (2020).

15. Singh, D. et al. Mechanisms of improved specificity of engineered Cas9s revealed by single-molecule FRET analysis. Nat. Struct. Mol. Biol. 25, 347–354 (2018).

16. Okafor, I. C. et al. Single molecule analysis of effects of non-canonical guide RNAs and specificity-enhancing mutations on Cas9-induced DNA unwinding. Nucleic Acids Res. 47, 11880–11888 (2019).

17. Bak, S. Y. et al. Quantitative assessment of engineered Cas9 variants for target specificity enhancement by single-molecule reaction pathway analysis. Nucleic Acids Res. 49, 11312–11322 (2021).

18. Wand, A. J. Bringing disorder and dynamics in protein allostery into focus. Proc. Natl. Acad. Sci. U. S. A. 114, 4278–4280 (2017).

19. Tzeng, S. R. & Kalodimos, C. G. Protein dynamics and allostery: An NMR view. Curr. Opin. Struct. Biol. 21, 62–67 (2011).

20. Wodak, S. J. et al. Allostery in Its Many Disguises: From Theory to Applications. Structure 27, 566–578 (2019).

21. East, K. W. et al. NMR and computational methods for molecular resolution of allosteric pathways in enzyme complexes. Biophys. Rev. 12, 155–174 (2020).

22. Liu, J. & Nussinov, R. Allostery: An Overview of Its History, Concepts, Methods, and Applications. PLOS Comput. Biol. 12, e1004966 (2016).

23. Guo, J. & Zhou, H. X. Protein Allostery and Conformational Dynamics. Chem. Rev. 116, 6503–6515 (2016).

24. Han, B., Liu, Y., Ginzinger, S. W. & Wishart, D. S. SHIFTX2: significantly improved protein chemical shift prediction. J. Biomol. NMR 50, 43–57 (2001).

25. Popovych, N., Sun, S., Ebright, R. H. & Kalodimos, C. G. Dynamically driven protein allostery. Nat. Struct. Mol. Biol. 13, 831–838 (2006).

26. Wand, A. J. The dark energy of proteins comes to light: conformational entropy and its role in protein function revealed by NMR relaxation. Curr. Opin. Struct. Biol. 23, 75–81 (2013).

27. Lisi, G. P. & Loria, J. P. Using NMR spectroscopy to elucidate the role of molecular motions in enzyme function. Prog. Nucl. Magn. Reson. Spectrosc. 92, 1–17 (2016).

28. Lisi, G. P. & Loria, J. P. Solution NMR Spectroscopy for the Study of Enzyme Allostery. Chem. Rev. 116, 6323–6369 (2016).

29. Ricci, C. G. et al. Deciphering Off-Target Effects in CRISPR-Cas9 through Accelerated Molecular Dynamics. ACS Cent. Sci. 5, 651–662 (2019).

30. Lindorff-Larsen, K. & Ferkinghoff-Borg, J. Similarity Measures for Protein Ensembles. PLoS One 4, e4203 (2009).

31. Schmid-Burgk, J. L. et al. Highly Parallel Profiling of Cas9 Variant Specificity. Mol. Cell 78, 794–800 (2020).

32. Sternberg, S. H., LaFrance, B., Kaplan, M. & Doudna, J. A. Conformational control of DNA target cleavage by CRISPR–Cas9. Nature 527, 110–113 (2015).

33. Delaglio, F. et al. NMRPipe: A Multidimensional Spectral Processing System Based on UNIX Pipes. J. Biomol. NMR 6, 277–293 (1995).

34. Lee, W., Tonelli, M. & Markley, J. L. NMRFAM-SPARKY: Enhanced software for biomolecular NMR spectroscopy. Bioinformatics 31, 1325–1327 (2015).

35. Skeens, E., East, K. W. & Lisi, G. P. 1H, 13C, 15 N backbone resonance assignment of the recognition lobe subdomain 3 (Rec3) from Streptococcus pyogenes CRISPR-Cas9. Biomol. NMR Assign. 15. 25–28 (2020).

36. Pervushin, K., Riek, R., Wider, G. & Wuthrich, K. Attenuated T2 Relaxation by Mutual Cancellation of Dipole-Dipole Coupling and Chemical Shift Anisotropy Indicates an Avenue to NMR Structures of Very Large Biological Macromolecules in Solution. Proc. Natl. Acad. Sci. U. S. A. 94, 12366–12371 (1997).

37. Grzesiek, S., Stahl, S. J., Wingfield, P. T. & Bax, A. The CD4 Determinant for Downregulation by HIV-1 Nef Directly Binds to Nef. Mapping of the Nef Binding Surface by NMR. Biochemistry 35, 10256–10261 (1996).

38. Zhu, G., Xia, Y., Nicholson, L. K. & Sze, K. H. Protein Dynamics Measurements by TROSY-Based NMR Experiments. J. Magn. Reson. 143, 423–426 (2000).

39. Bieri, M., d’Auvergne, E. J. & Gooley, P. R. relaxGUI: A New Software for Fast and Simple NMR Relaxation Data Analysis and Calculation of ps-ns and us Motion of Proteins. J. Biomol. NMR 50, 147–155 (2011).

40. Lipari, G. & Szabo, A. Model-Free Approach to the Interpretation of Nuclear Magnetic Resonance Relaxation in Macromolecules. 1. Theory and Range of Validity. J. Am. Chem. Soc. 104, 4546–4559 (1982).

41. Loria, J. P., Rance, M. & Palmer 3rd, A. G. A Relaxation-Compensated Carr-Purcell-Meiboom-Gill Sequence for Characterizing Chemical Exchange by NMR Spectroscopy. J. Am. Chem. Soc. 121, 2331–2332 (1999).

42. Bieri, M. & Gooley, P. R. Automated NMR relaxation dispersion data analysis using NESSY. BMC Bioinformatics 12, 421 (2011).

43. Luz, Z. & Meiboom, S. Nuclear magnetic resonance study of the protolysis of trimethylammonium ion in aqueous solution-order of the reaction with respect to solvent. J. Chem. Phys. 39, 366–370 (1963).

44. Carver, J. P. & Richards, R. E. A general two-site solution for the chemical exchange produced dependence of T2 upon the carr-Purcell pulse separation. J. Magn. Reson. 6, 89–105 (1972).

45. Kabsch, W. XDS. Acta Crystallogr. D Biol. Cristallogr. 66, 125–132 (2010).

46. Winn, M. D. et al. Overview of the CCP4 Suite and its Current Developments. Acta Crystallogr. D Biol. Cristallogr. 67, 235–242 (2011).

47. Liebschner, D. et al. Macromolecular structure determination using X-rays, neutrons and electrons: recent developments in *Phenix*. Acta Crystallogr. D Struct. Biol. 75, 861–877 (2019).

48. Emsley, P., Lohkamp, B., Scott, W.G. & Cowtan, K. Features and Development of Coot. Acta Crystallogr. D Biol. Crystallogr. 66, 486–501 (2010).

49. Yu, L., Li, D. W. & Brüschweiler, R. Balanced Amino-Acid-Specific Molecular Dynamics Force Field for the Realistic Simulation of Both Folded and Disordered Proteins. J. Chem. Theory Comput. 16, 1311–1318 (2020).

50. Nierzwicki, L. et al. Enhanced specificity mutations perturb allosteric signaling in CRISPR-Cas9. Elife 10, e73601 (2021).

51. Ivani, I. et al. Parmbsc1: a refined force field for DNA simulations. Nat. Methods 13, 55– 58 (2016).

52. Banas, P. et al. Performance of Molecular Mechanics Force Fields for RNA Simulations: Stability of UUCG and GNRA Hairpins. J. Chem. Theor. Comput. 6, 3836–3849 (2010).

53. Zgarbova, M. et al. Refinement of the Cornell et al. Nucleic Acids Force Field Based on Reference Quantum Chemical Calculations of Glycosidic Torsion Profiles. J. Chem. Theory Comput. 7, 2886–2902 (2011).

54. Jorgensen, W. L., Chandrasekhar, J., Madura, J. D., Impey, R. W. & Klein, M. L. Comparison of simple potential functions for simulating liquid water. J. Chem. Phys. 79, 926–935 (1983).

55. Turq, P., Lantelme, F. & Friedman, H. L. Brownian Dynamics: Its Applications to Ionic Solutions. J. Chem. Phys. 66, 3039 (1977).

56. Berendsen, H. J. C., Postma, J. P. M., van Gunsteren, W. F., DiNola, A. & Haak, J. R. Molecular Dynamics with Coupling to an External Bath. J. Chem. Phys. 81, 3684 (1984).

57. Palermo, G. et al. Protospacer Adjacent Motif-Induced Allostery Activates CRISPR-Cas9. J. Am. Chem. Soc. 139, 16028–16031 (2017).

58. Case, D. A. et al. AMBER 2020. Univ. of California, San Francisco (2020).

59. Vögele, M., et al. Systematic Analysis of Biomolecular Conformational Ensembles with PENSA. Preprint at 10.48550/arXiv.2212.02714 (2022).

60. Sethi, A., Eargle, J., Black, A. A. & Luthey-Schulten, Z. Dynamical networks in tRNA: protein complexes. Proc. Natl. Acad. Sci. U. S. A. 106, 6620–6625 (2009).

61. Arantes, P. R., Patel, A. C. & Palermo, G. Emerging Methods and Applications to Decrypt Allostery in Proteins and Nucleic Acids. J. Mol. Biol. 434, 167518 (2022)

62. Doshi, U., Holliday, M. J., Eisenmesser, E. Z. & Hamelberg, D. Dynamical network of residue-residue contacts reveals coupled allosteric effects in recognition, catalysis, and mutation. Proc. Natl. Acad. Sci. U. S. A. 113, 4735–40 (2016).

63. Dodd, T. et al. Polymerization and editing modes of a high-fidelity DNA polymerase are linked by a well-defined path. Nat. Commun. 11, 5379 (2020).

64. Melo, M. C. R., Bernardi, R. C., de la Fuente-Nunez, C. & Luthey-Schulten, Z. Generalized correlation-based dynamical network analysis: a new high-performance approach for identifying allosteric communications in molecular dynamics trajectories. J. Chem. Phys. 153, 134104 (2020).

65. Lange, O. F. & Grubmüller, H. Generalized correlation for biomolecular dynamics. Proteins Struct. Funct. Genet. 62, 1053–1061 (2006).

66. Floyd, R. W. Algorithm 97: Shortest path. Commun. ACM 5, 345 (1962).

67. Yen, J. Y. Finding the *K* Shortest Loopless Paths in a Network. Manage. Sci. 17, 712–716 (1971).

68. Hagberg, A., Schult, D. & Swart, P. Exploring Network Structure, Dynamics, and Function using NetworkX in Proceedings of the 7th Python in Science conference (SciPy 2008). G Varoquaux, T Vaught, J Millman (Eds.), pp. 11-15.

