## Supporting Information for "High-Fidelity, Hyper-Accurate, and Evolved Mutants Rewire Atomic Level Communication in CRISPR-Cas9"

### Table of Contents

**Supplementary Materials and Methods**

Generalized Correlation (GC) analysis

Contact analysis

**Supplementary Figures**

**Supplementary Fig. 1.** Secondary structure and thermal stability of WT Rec3 and the Rec3 variants

**Supplementary Fig. 2.** $^{1}$H-$^{15}$N TROSY HSQC spectrum of the WT Rec3 domain

**Supplementary Fig. 3.** Fast timescale dynamics (ps – ns) of the WT Rec3 domain by NMR

**Supplementary Fig. 4.** $^{1}$H$^{15}$N TROSY HSQC spectral overlay of WT Rec3 and the Rec3 variants

**Supplementary Fig. 5.** $^{1}$H-$^{13}$C spectral overlays of WT Rec3 and the Rec3 variants

**Supplementary Fig. 6.** Carbon chemical shift perturbations of the Rec3 variants

**Supplementary Fig. 7.** Longitudinal ($R_1$) relaxation rates for the Rec3 variants

**Supplementary Fig. 8.** Longitudinal ($R_2$) relaxation rates for the Rec3 variants

**Supplementary Fig. 9.** $^1$H-$^{15}$N NOE measurements for the Rec3 variants

**Supplementary Fig. 10.** Order parameters ($S^2$) for the Rec3 variants

**Supplementary Fig. 11.** CPMG relaxation dispersion for WT Rec3 and the Rec3 variants

**Supplementary Fig. 12.** Principal Component Analysis (PCA)

**Supplementary Fig. 13.** Analysis of allosteric pathways connecting the Rec3 α-helix 37 to the HNH catalytic core in the WTCas9 and its HF1, Hypa, and Evo variants.

**Supplementary Fig. 14.** Analysis of allosteric pathways connecting the Rec3 mutation sites to the HNH catalytic core in the WT and Cas9-HF1

**Supplementary Fig. 15.** Analysis of allosteric pathways connecting the Rec3 mutation sites to the HNH catalytic core in the WT and evoCas9

**Supplementary Tables**

**Table S1.** Conformational exchange parameters for residues in WT Rec3 via CPMG relaxation dispersion experiments.

**Table S2.** Conformational exchange parameters for residues in HF1 Rec3 via CPMG relaxation dispersion experiments.

**Table S3.** Conformational exchange parameters for residues in Hypa Rec3 via CPMG relaxation dispersion experiments.

**Table S4.** Conformational exchange parameters for residues in Evo Rec3 via CPMG relaxation dispersion experiments

**Supplementary References**

### Supplementary Materials and Methods

#### Generalized Correlation (GC) analysis

Generalized Correlation ($GC$) anaysis[1] was used to quantify the systems' correlations based on Mutual Information ($MI$) and to build dynamical network models of Cas9 and its variants. Through this analysis, correlations between a pair of nodes are described independently on their relative orientation, capturing non–linear contributions to correlations. In this analysis, two variables ($x_i$, $x_j$) can be considered correlated when their joint probability distribution, $p(x_i,\, x_j)$, is smaller than the product of their marginal distributions, $p(x_i) \cdot p(x_j)$. The $MI$ is a measure of the degree of correlation between $x_i$ and $x_j$ defined as a function of $p(x_i,\, x_j)$ and $p(x_i) \cdot p(x_j)$ according to:

$$MI\,[x_i,\, x_j] = \iint p(x_i,\, x_j)\, ln \frac{p(x_i, x_j)}{p(x_i) \cdot p(x_j)} dx_i dx_j \tag{1}$$

The $MI$ is closely related to the definition of the Shannon entropy, $H[x]$, i.e., the expectation value of a random variable $x$, having a probability distribution $p(x_i)$

$$H[x] = -\int p(x)\, ln\, p(x) dx \tag{2}$$

and it can be computed as:

$$MI\,[x_i,\, x_j] = H\,[x_i] + H\,[x_j] - H\,[x_i, x_j] \tag{3}$$

where $H\,[x_i]$ and $H\,[x_j]$ are the marginal Shannon entropies, and $H\,[x_i, x_j]$ is the joint entropy. Since $MI$ varies from 0 to +∞, normalized generalized correlation ($GC$) coefficients, ranging from 0 (independent variables) to 1 (fully correlated variables), are defined as:

$$GC\,[x_i,\, x_j] = \left\{1 - e^{-\frac{2MI[x_i, x_j]}{d}}\right\}^{-\frac{1}{2}} \tag{4}$$

where $d = 3$ is the dimensionality of $x_i$ and $x_j$.

In the present work, $GCs$ were computed using the recent high-performance $GC$-based dynamical network analysis tool by Luthey-Schulten.[2] In a second phase, the $GCs$ were used as a metric to build dynamical network models of the systems as described in the main text.

**Contact analysis**

To describe the interactions between the RNA:DNA hybrid and the HNH domain in the WTCas9 and in its three variants, we conducted an in-depth contact analysis. For each system, we first computed the interfacial contact maps considering contacts formed at a distance cut-off of 4.5 Å between any heavy atom. Then, the frequency of each formed contact, $f$, was computed as the ratio between the number of frames each contact is formed and the total number of frames. Finally, to detail the gain or loss of contact stability moving from one system to another (i.e., WTCas9 *vs.* Cas9-HF1; HypaCas9 or evoCas9), we computed the differential contact stability. The difference in stability of contacts ($\Delta f_{A-B}$) in systems $A$ and $B$, is computed as $\Delta f_{A-B} = f_A - f_B$, where the stability of contacts in systems $A$ and $B$ is represented by their frequencies, $f_A$ and $f_B$, respectively. As $f$ varies from 0 (contacts never formed) to 1 (contacts accounted in all frames), $\Delta f_{A-B}$ varies from -1 to +1, where a $\Delta f_{A-B} < 0$ corresponds to contacts relatively more stable in system $B$ and $\Delta f_{A-B} > 0$ corresponds to contacts relatively more stable in system $A$. Differential contact maps were plotted applying a cut-off of $|\Delta f_{A-B}| \geq 0.1$, which accounts for contacts that are more stable in one system (e.g., $A$) for more than 10 % with respect to the other (e.g., $B$).

### Supplementary Figures

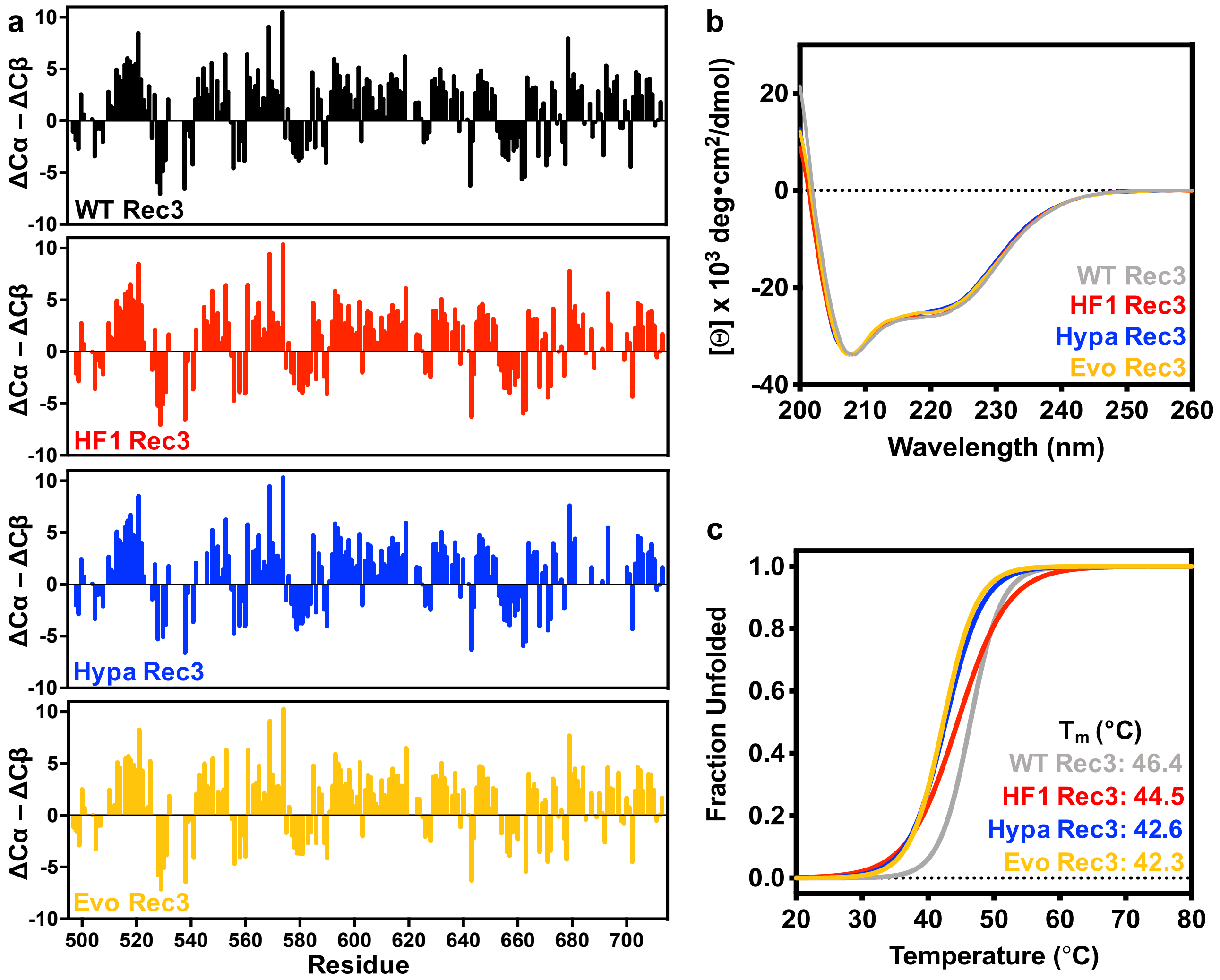


**Supplementary Fig. 1. Secondary structure and thermal stability of WT Rec3 and the Rec3 variants. a.** Secondary structure of WT Rec3 (black), HF1 Rec3 (red), Hypa Rec3 (blue), and Evo Rec3 (yellow) are estimated from NMR-derived Cα and Cβ chemical shifts and calculated relative to random coil reference values for each residue (ΔCα-ΔCβ). Positive values correspond to alpha-helical secondary structure, and negative values denote beta-sheet secondary structure. **b.** Far-UV circular dichroism spectroscopy of WT Rec3 (grey), HF1 Rec3 (red), Hypa Rec3 (blue), and Evo Rec3 (yellow) at 25°C. **c.** Thermal denaturation profiles of WT Rec3 and the Rec3 variants (λ = 208 nm), normalized as a fraction of unfolded protein.

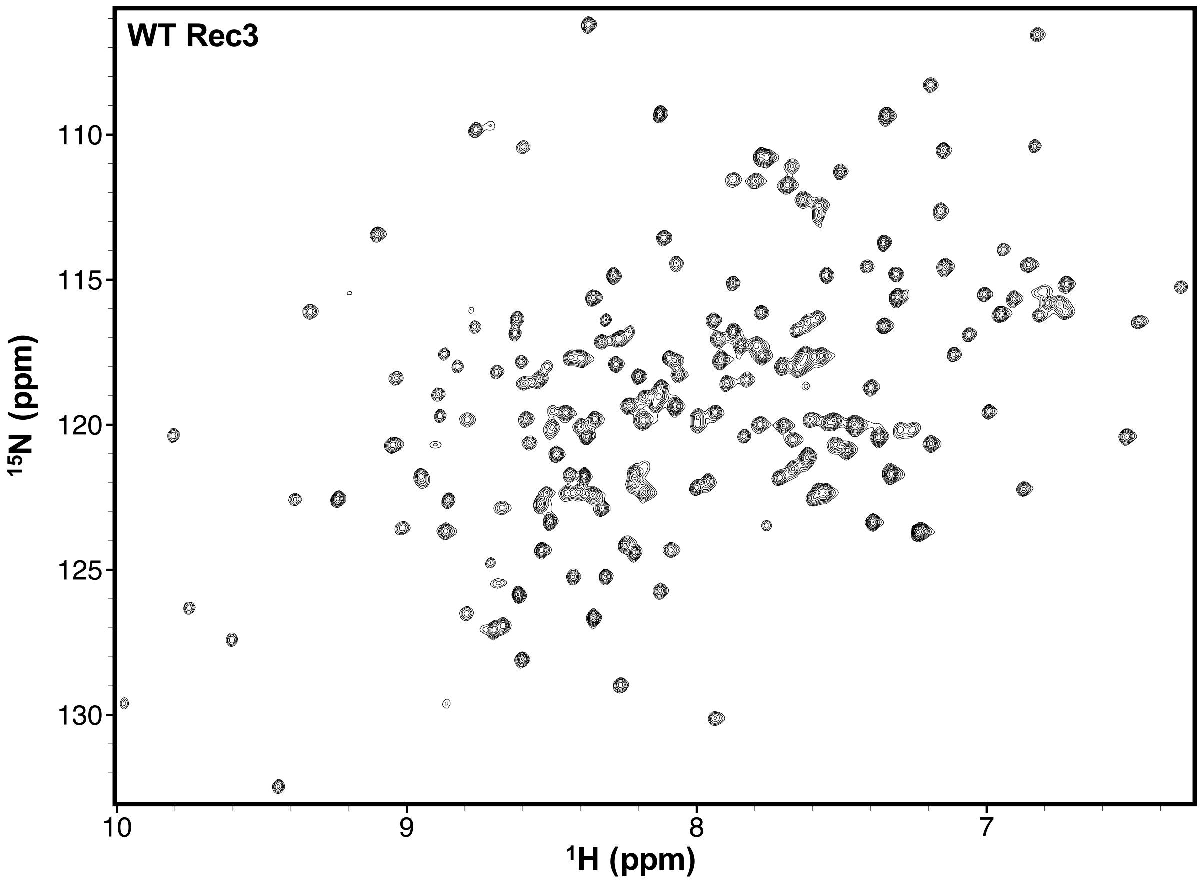


**Supplementary Fig. 2**. **$^{1}$H-$^{15}$N TROSY HSQC spectrum of the WT Rec3 domain.** The WT Rec3 spectrum was acquired on a 650 μM sample at 30°C and 600 MHz. Backbone resonance assignments can be found in the BMRB (#50389).

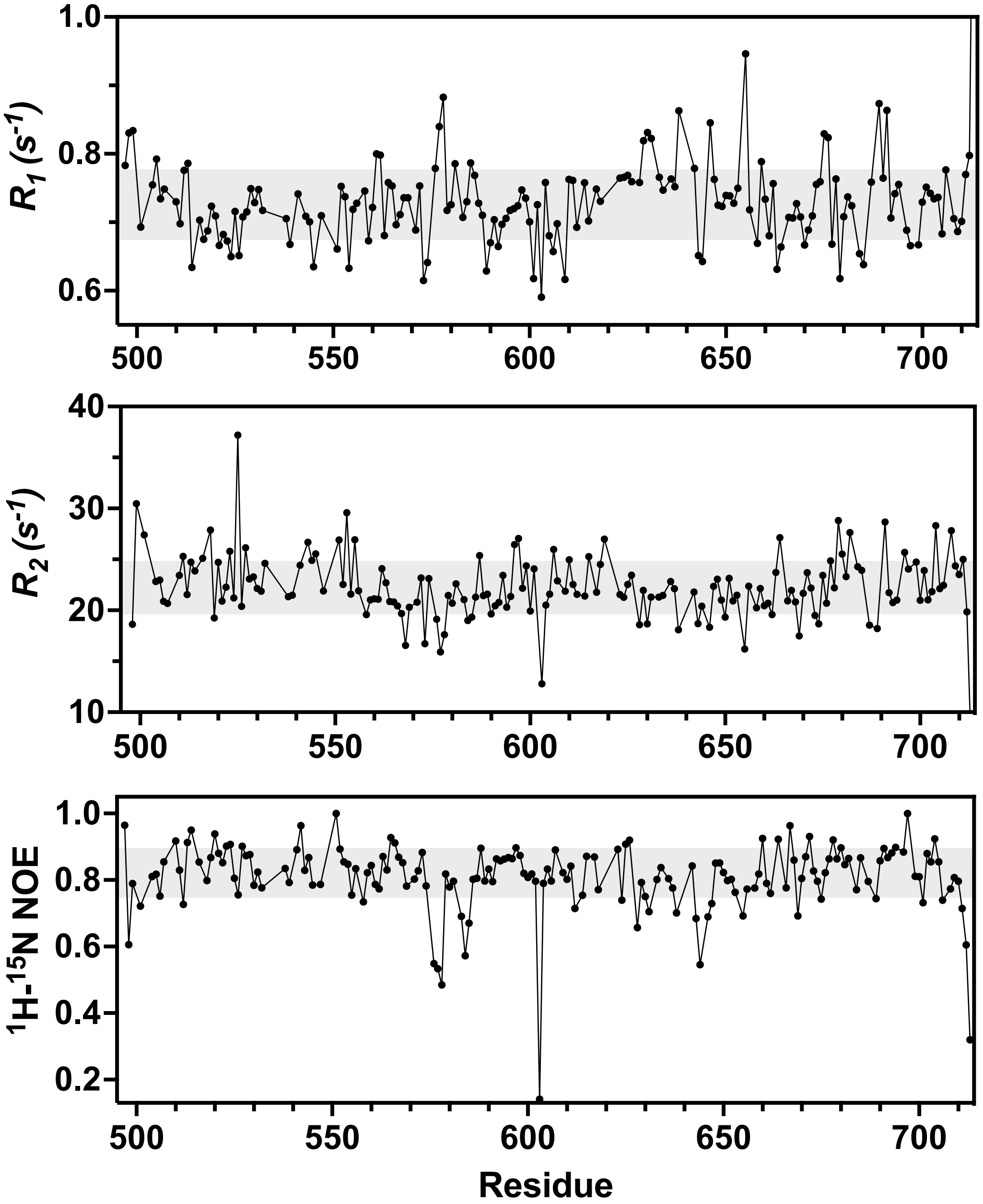


**Supplementary Fig. 3. Fast timescale dynamics (ps – ns) of the WT Rec3 domain by NMR.** Plots of longitudinal ($R_1$) and transverse ($R_2$) relaxation rates and $^1$H-[$^{15}$N] NOE values for WT Rec3. Gray shaded bars denote ±1.5σ of the 10% trimmed mean for each parameter.

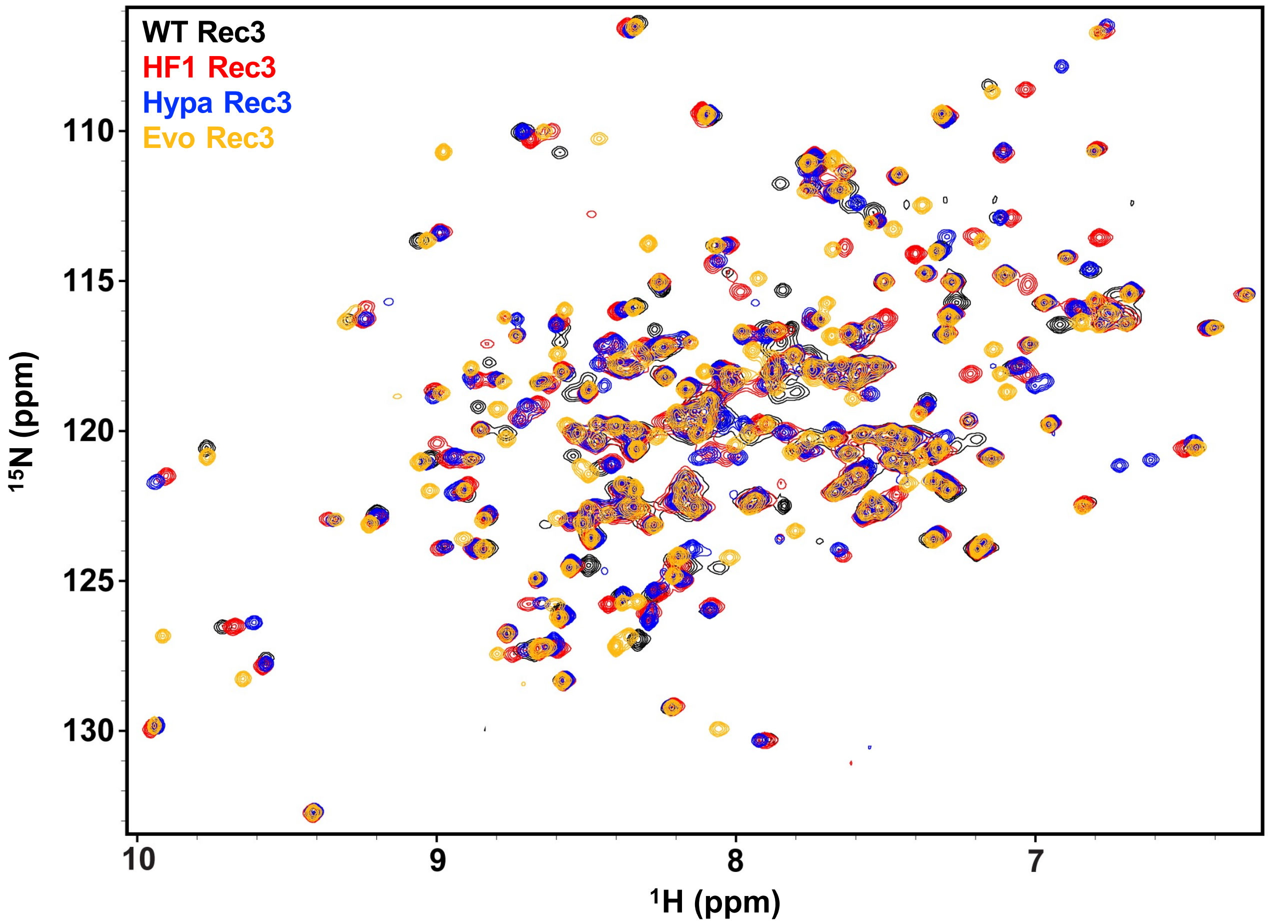


**Supplementary Fig. 4. $^{1}H^{15}N$ TROSY HSQC spectral overlay of WT Rec3 and the Rec3 variants.** Spectra were acquired on 650 μM samples at 30°C and 600 MHz for WT Rec3 (black), HF1 Rec3 (red), Hypa Rec3 (blue), and Evo Rec3 (yellow).

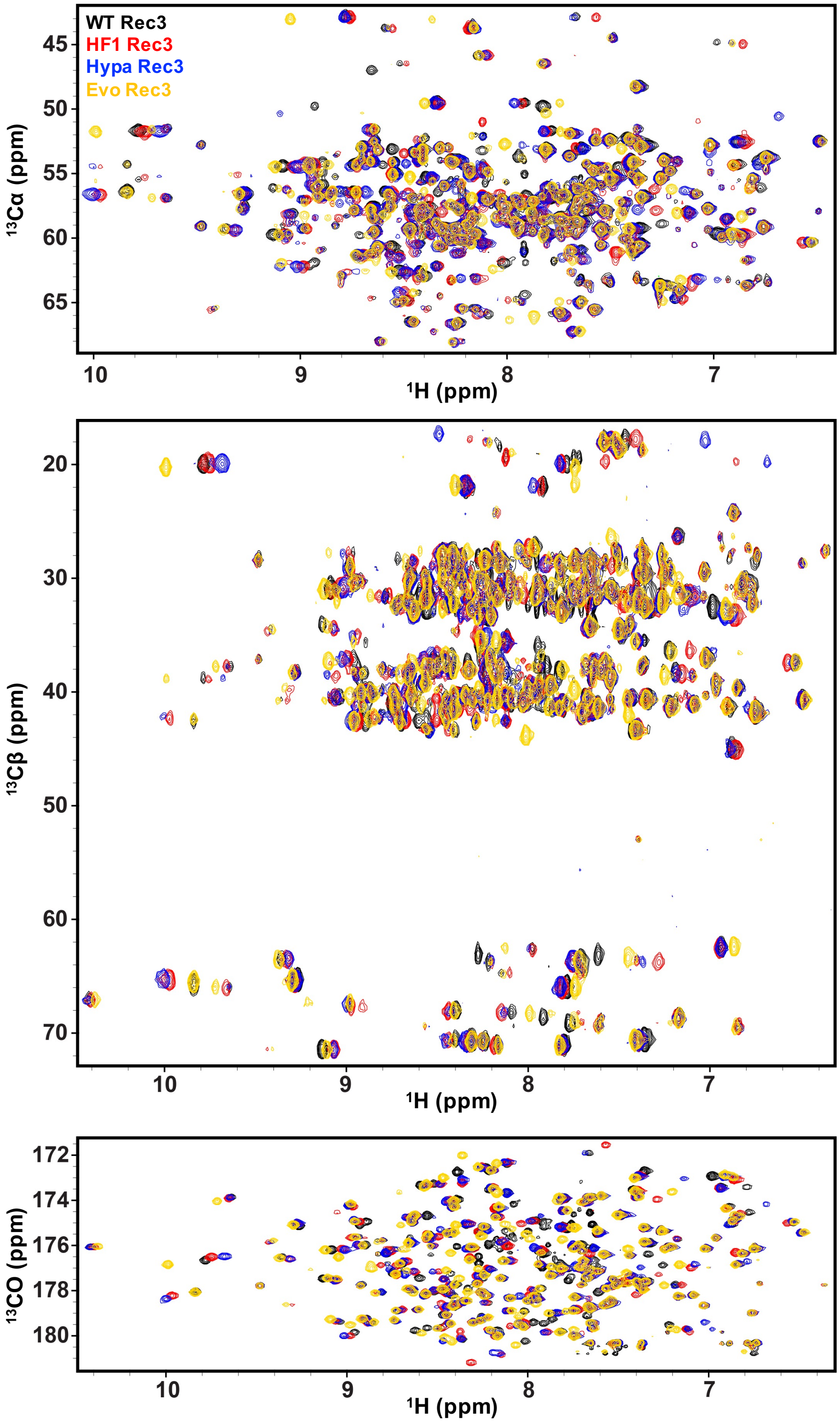


**Supplementary Fig. 5. $^{1}$H-$^{13}$C spectral overlays of WT Rec3 and the Rec3 variants.** $^{13}$Cα (top), $^{13}$Cβ (middle), and $^{13}C_O$ (bottom) spectra were acquired on 650 μM samples at 30°C and 600 MHz for WT Rec3 (black), HF1 Rec3 (red), Hypa Rec3 (blue), and Evo Rec3 (yellow).

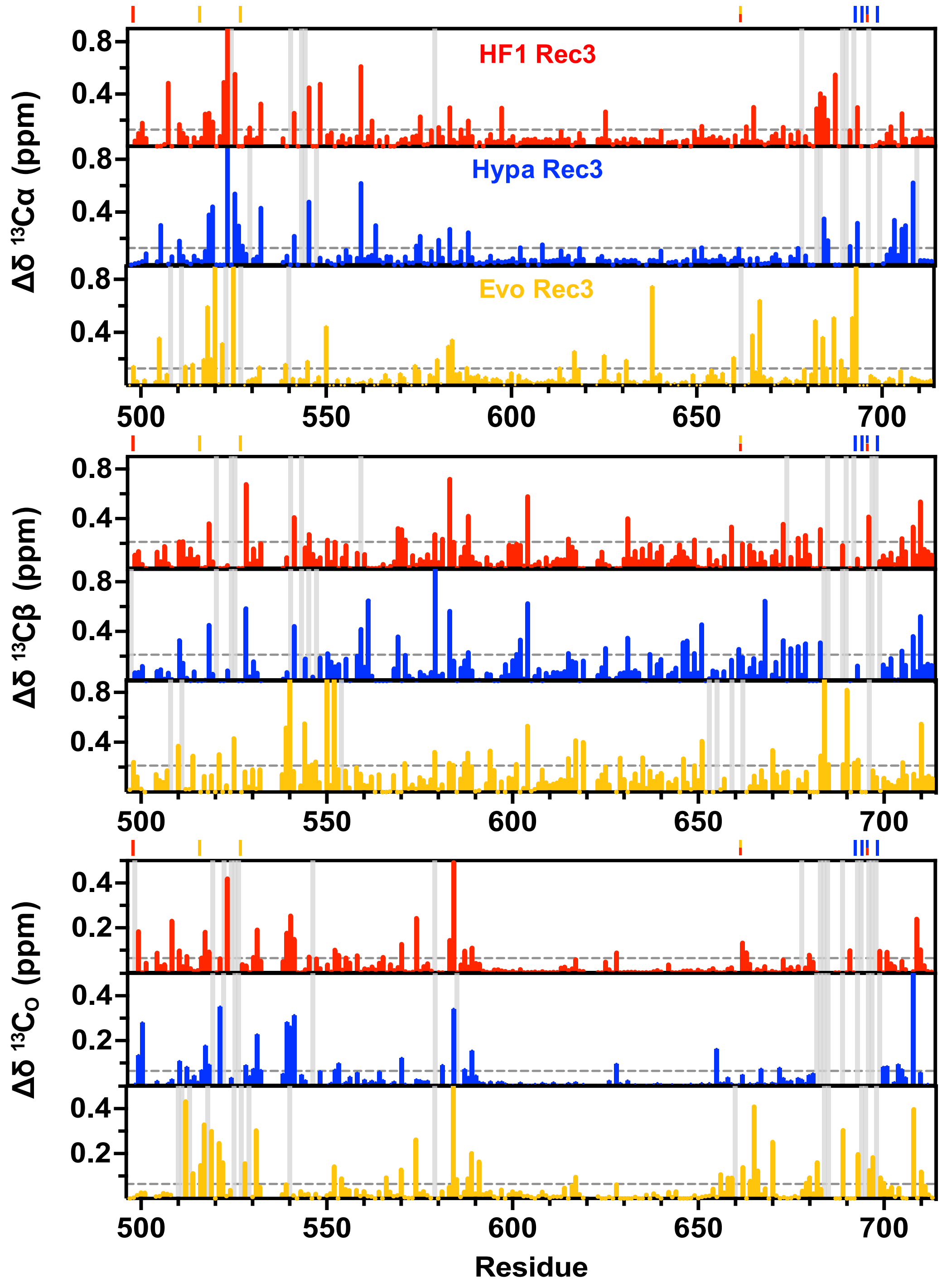


**Supplementary Fig. 6. Carbon chemical shift perturbations of the Rec3 variants.** $^{13}$Cα (top), $^{13}$Cβ (middle), and $^{13}$C$_O$ (bottom) chemical shifts perturbations (Δδ) are plotted for HF1 Rec3 (red), Hypa Rec3 (blue), and Evo Rec3 (yellow) relative to WT Rec3. Sites of mutation are indicated by coloured lines above the plots, grey vertical bars represent line-broadened resonances, and grey dashed horizontal lines denote 1.5σ above the 10% trimmed mean of all shifts for each carbon position.

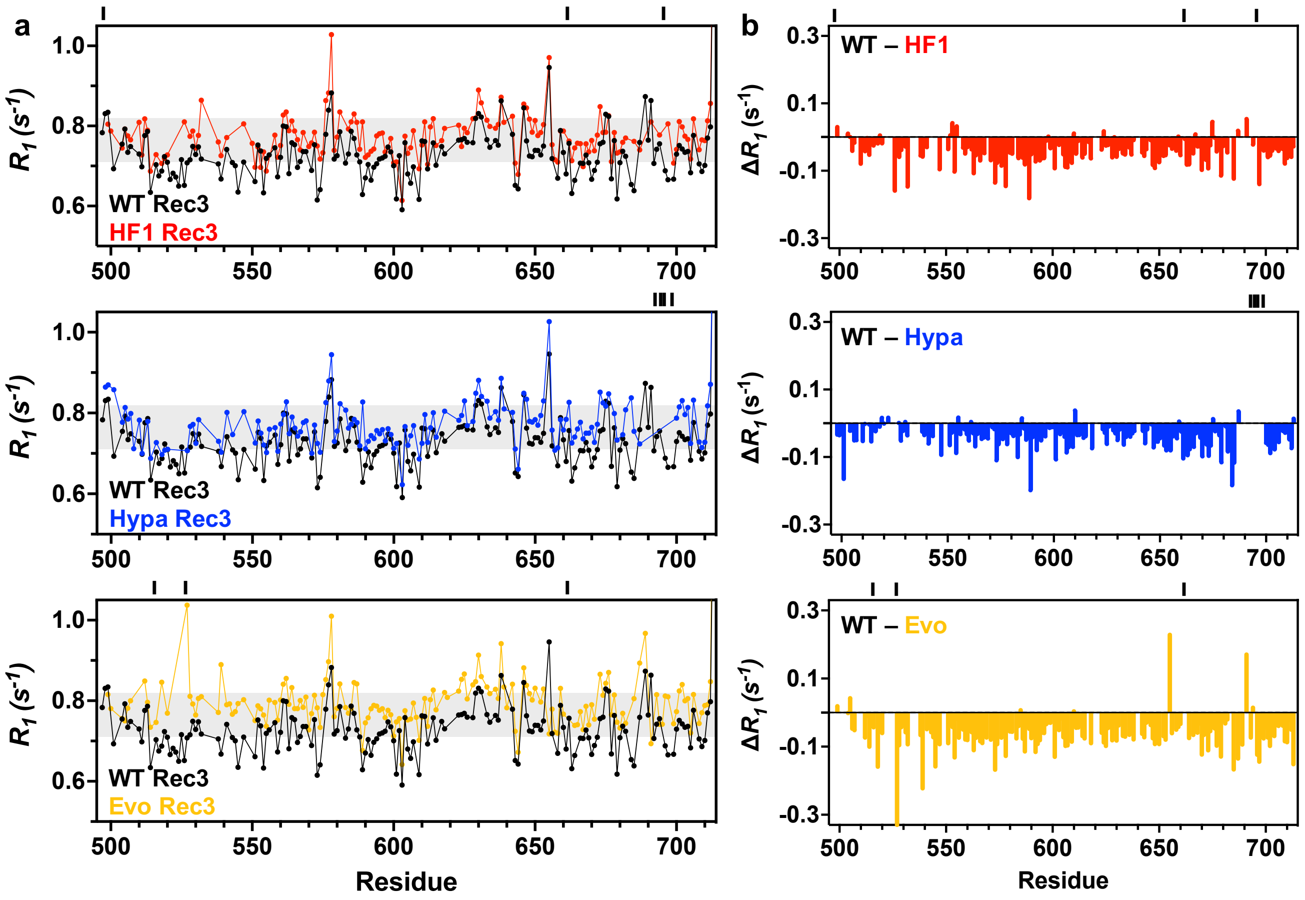


**Supplementary Fig. 7. Longitudinal ($R_1$) relaxation rates for the Rec3 variants. a.** Plots of $R_1$ relaxation rates for HF1 Rec3 (red), Hypa Rec3 (blue), and Evo Rec3 (yellow) are overlaid with WT Rec3 (black) for comparison. The grey shaded area denotes ±1.5σ from the 10% trimmed mean of all data sets. **b.** $\Delta R_1$ plots highlighting changes in $R_1$ relaxation rates between WT Rec3 and the Rec3 variants.

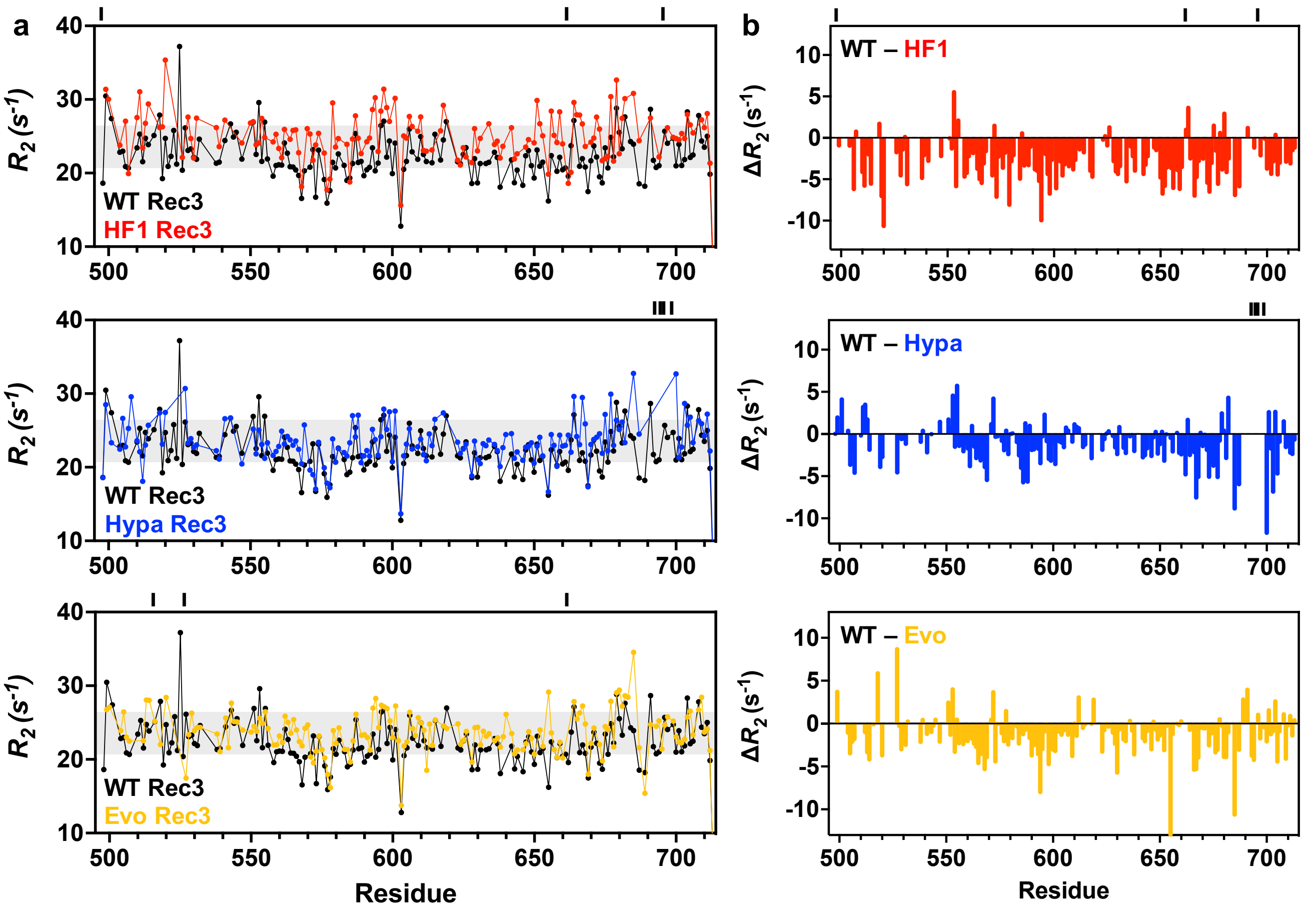


**Supplementary Fig. 8. Longitudinal ($R_2$) relaxation rates for the Rec3 variants. a.** Plots of $R_2$ relaxation rates for HF1 Rec3 (red), Hypa Rec3 (blue), and Evo Rec3 (yellow) are overlaid with WT Rec3 (black) for comparison. The grey shaded areas denote ±1.5σ from the 10% trimmed mean of all data sets. **b.** $\Delta R_2$ plots highlighting changes in $R_1$ relaxation rates between WT Rec3 and the Rec3 variants.

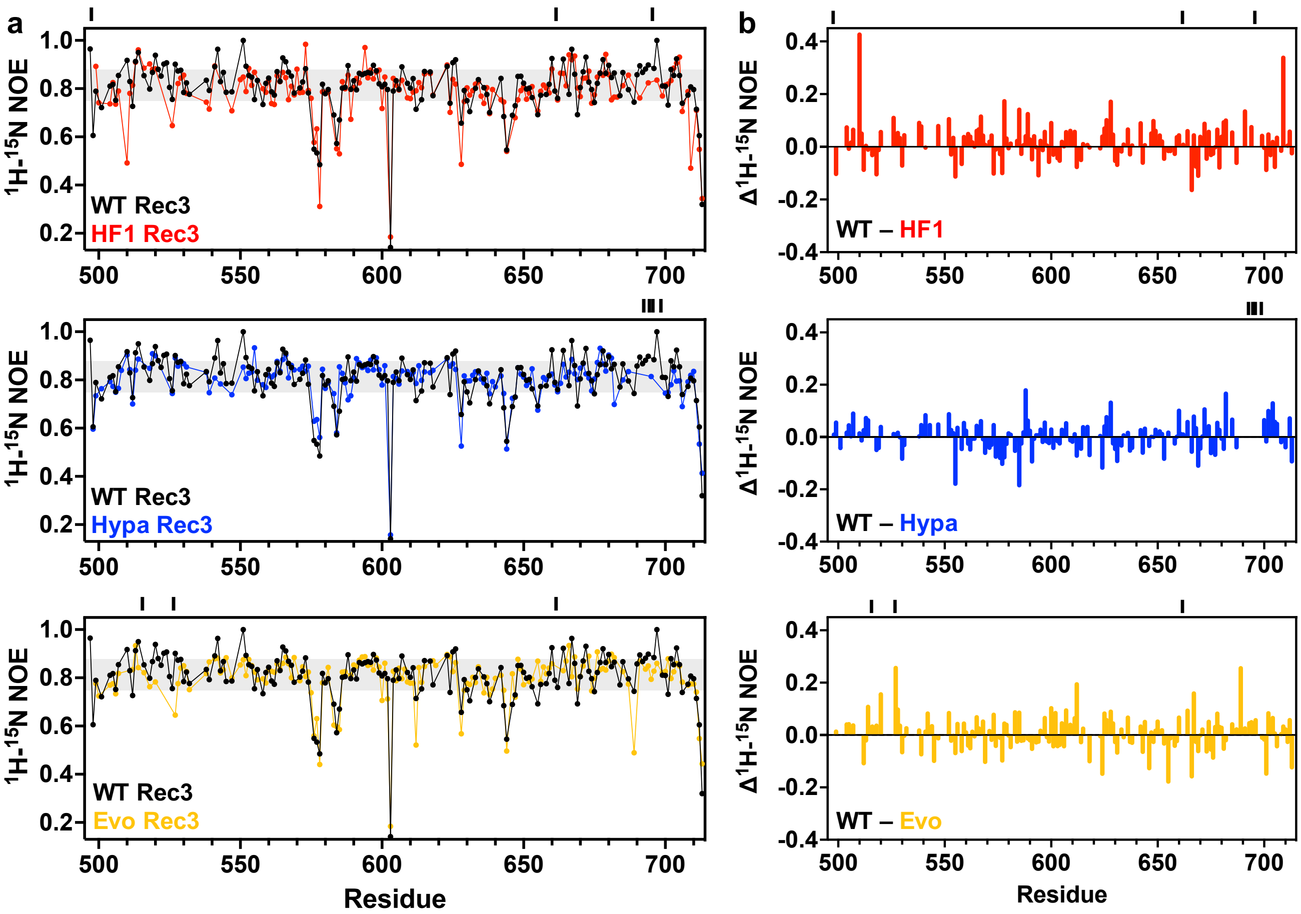


**Supplementary Fig. 9. $^{1}H$-$^{15}N$ NOE measurements for the Rec3 variants. a.** Plots of $^{1}H$-$^{15}N$ NOE values for HF1 Rec3 (red), Hypa Rec3 (blue), and Evo Rec3 (yellow) are overlaid with WT Rec3 (black) for comparison. The grey shaded area denotes ±1.5σ from the 10% trimmed mean of all data sets. **b.** $\Delta^{1}H$-$^{15}N$ NOE plots highlighting changes in $^{1}H$-$^{15}N$ NOE values between WT Rec3 and the Rec3 variants.

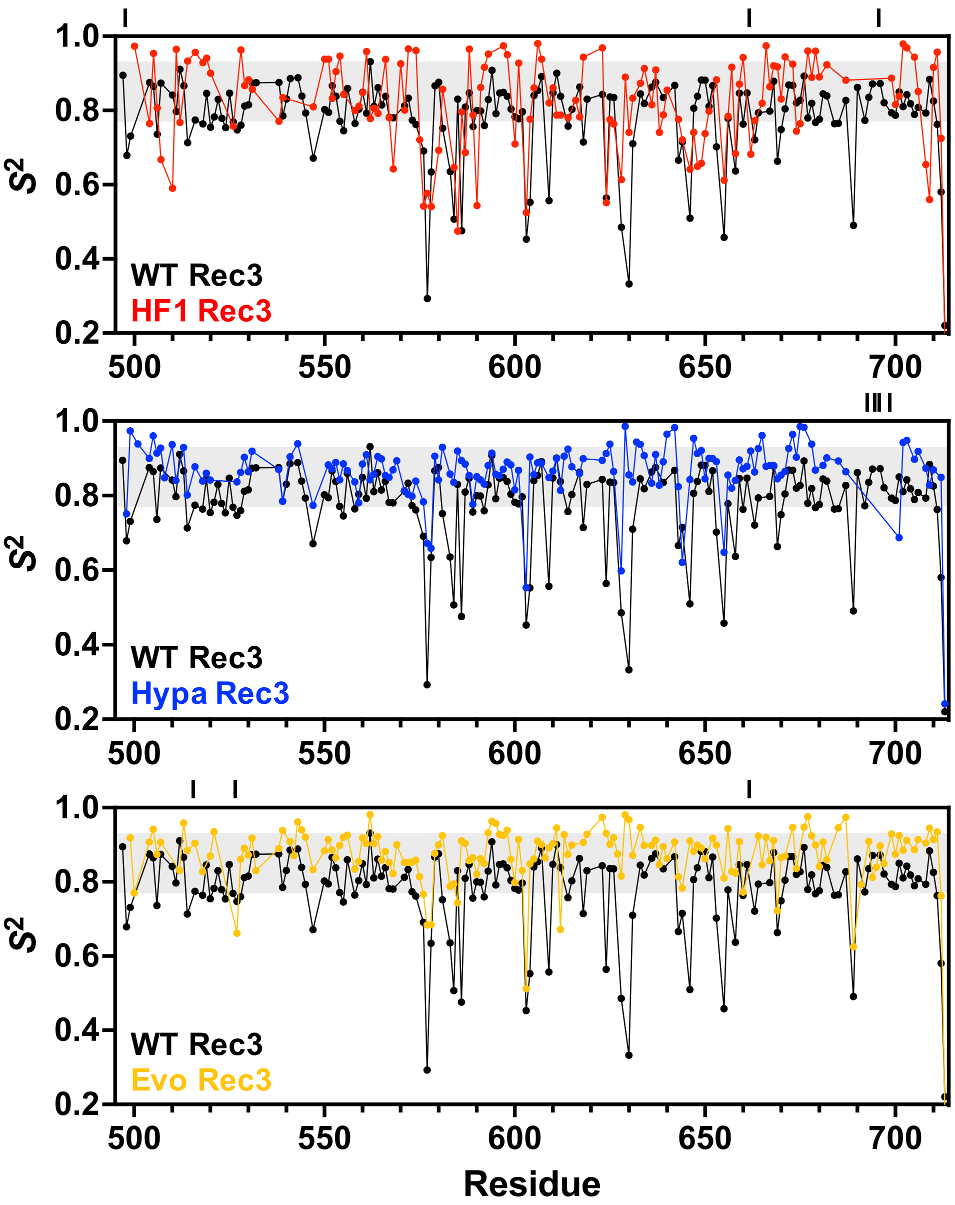


**Supplementary Fig. 10. Order parameters ($S^2$) for the Rec3 variants.** Plots of $S^2$ values for HF1 Rec3 (red), Hypa Rec3 (blue), and Evo Rec3 (yellow) are overlaid with WT Rec3 (black) for comparison. The grey shaded area denotes ±1.5σ from the 10% trimmed mean of all data sets.

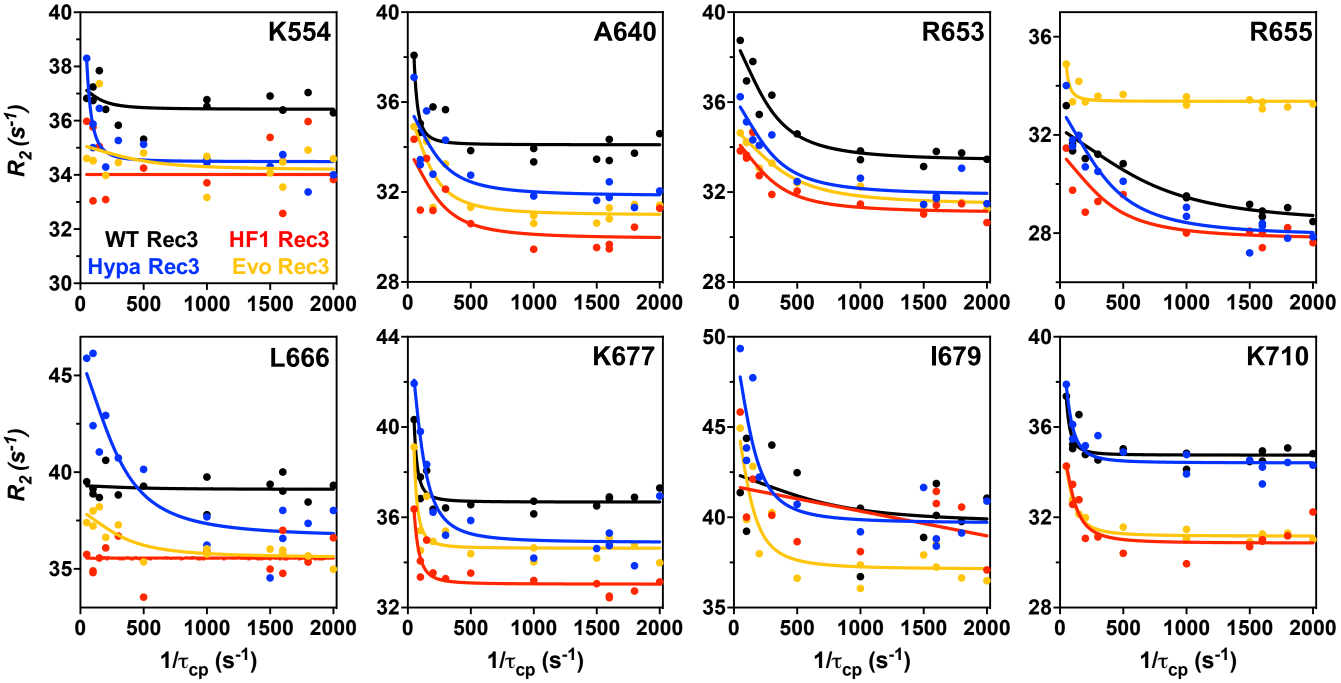


**Supplementary Fig. 11. CPMG relaxation dispersion for WT Rec3 and the Rec3 variants.** Dispersion profiles for selected residues in WT Rec3 (black), HF1 Rec3 (red), Hypa Rec3 (blue), and Evo Rec3 (yellow) are shown. Some of the representative plots show residues where μs – ms motions are conserved in WT Rec3 and the three variants (A640, R653, K677, K710), while other plots show gain (curved profiles) or loss (flat profiles) of μs – ms motions in one or more of the variants (K554, R655, L666, I679).

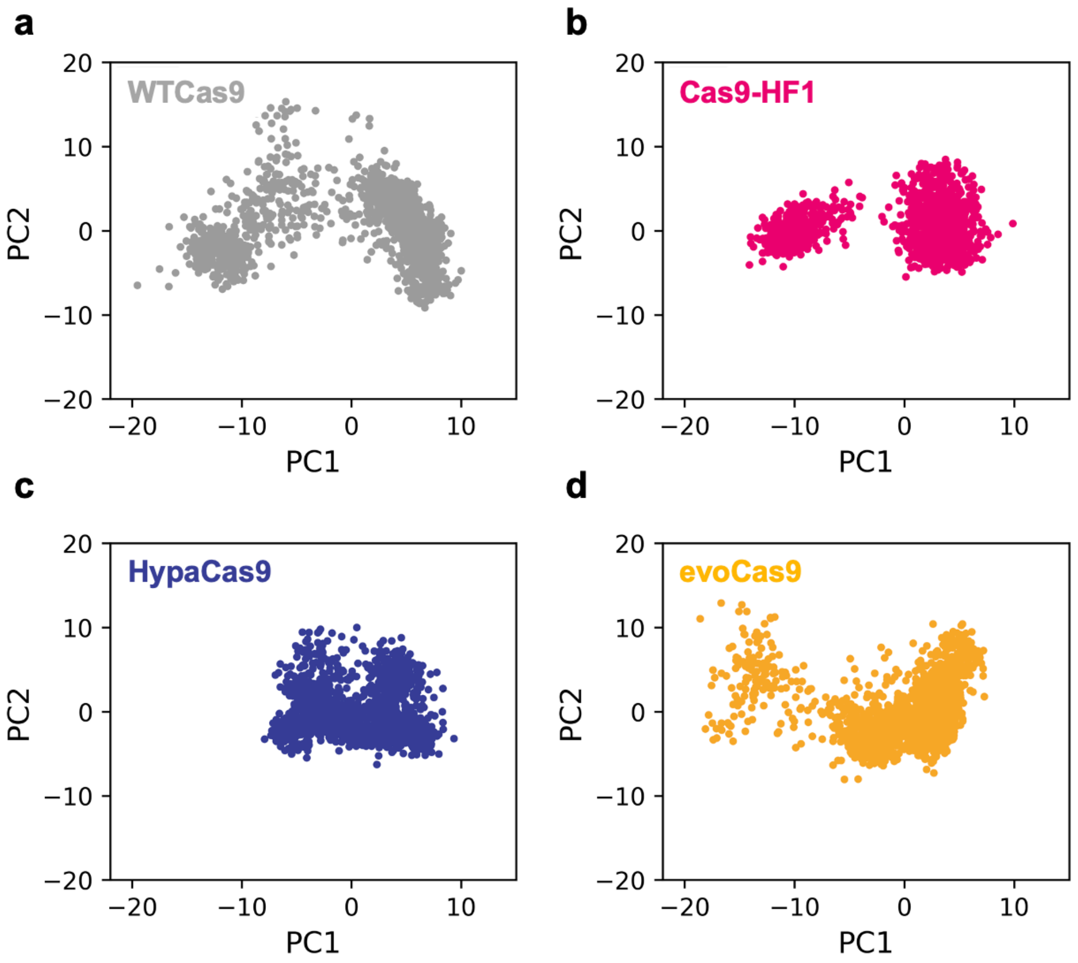


**Supplementary Fig. 12. Principal Component Analysis (PCA).** Projections of the first principal and second principal components (PC1 *vs.* PC2), derived from PCA of the HNH domains in (**a**) WT, (**b**) Cas9-HF1, (**c**) HypaCas9, and (**d**) evoCas9 complexes. PCA was performed on the backbone Cα atoms. The collective ensembles were combined and subjected to root mean square (RMS)-fit and alignment to the same reference configuration, removing rotational and translational motions, to ensure a consistent eigenbasis on all compared systems.

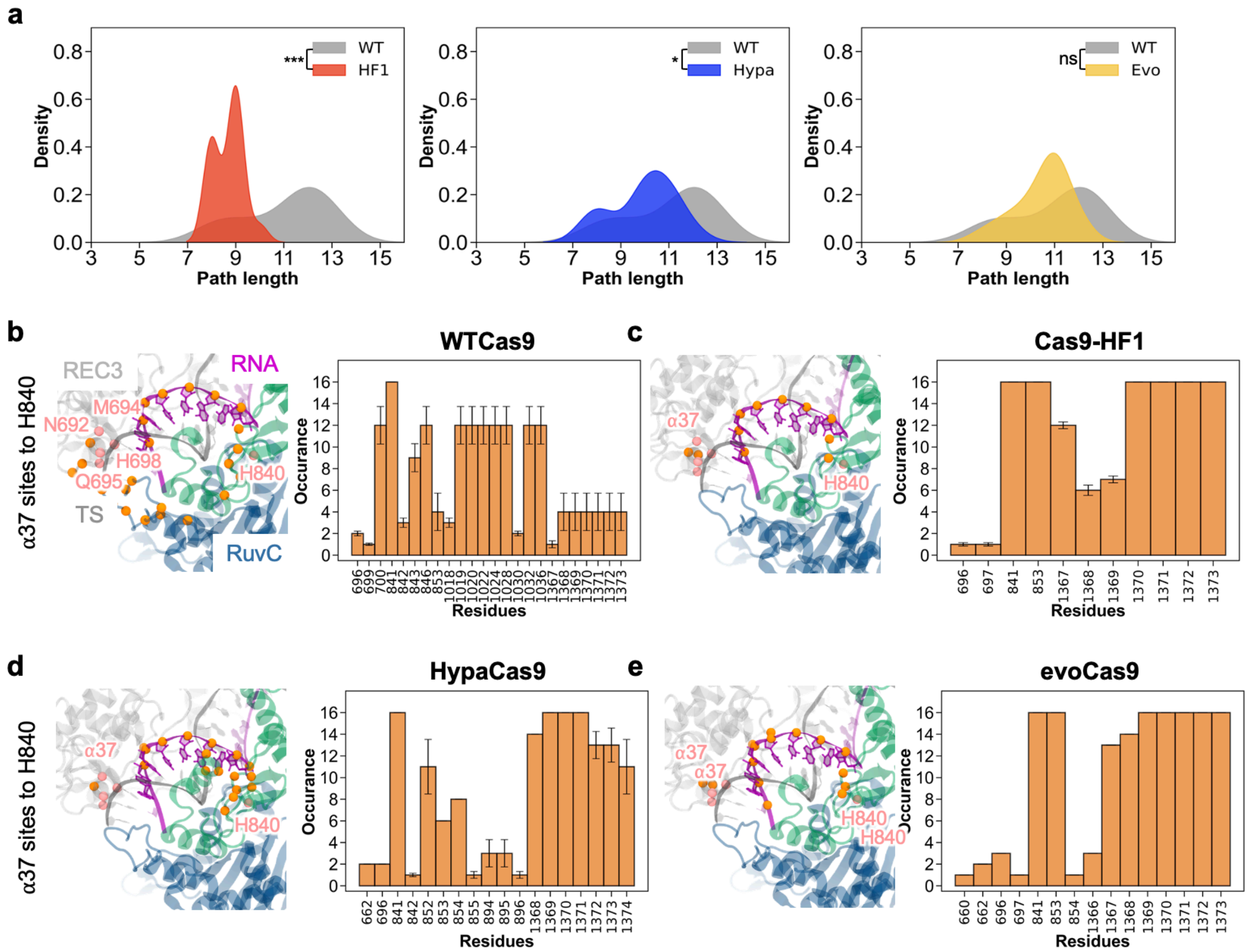

**Supplementary Fig. 13. Analysis of allosteric pathways connecting the Rec3 α-helix 37 to the HNH catalytic core in the WTCas9 and its HF1, Hypa, and Evo variants. a.** Kernel density estimation of pathlengths in terms of number edges. Pathways are computed connecting the Rec3 mutation sites clustered in the α-helix 37 (residues 692, 694, 695, and 698) and the HNH catalytic residue H840. The HF1 (red), Hypa (blue), and Evo (yellow) variants are compared to the WT (gray) Cas9. The distributions of path lengths were plotted by combining all pathways connecting the α37 mutation sites to H840. The statistical significance among the distributions was calculated using a two-tailed unpaired t-test (P-value reference: not significant, ns P > 0.05, * P ≤ 0.05, ** P ≤ 0.005, *** P ≤ 0.0005). **b-c.** Occurrence of residues in the optimal and top five suboptimal pathways communicating the Rec3 mutation sites to the HNH catalytic site in the WT (**b**) and Hypa (**c**) Cas9. Error bars correspond to the standard error of the mean computed over three sample pools obtained from ~16μs of aggregated sampling for each of the systems. Signalling pathways are also shown on the three-dimensional structure using spheres of different colours: pink (source and sink) and orange (path nodes).

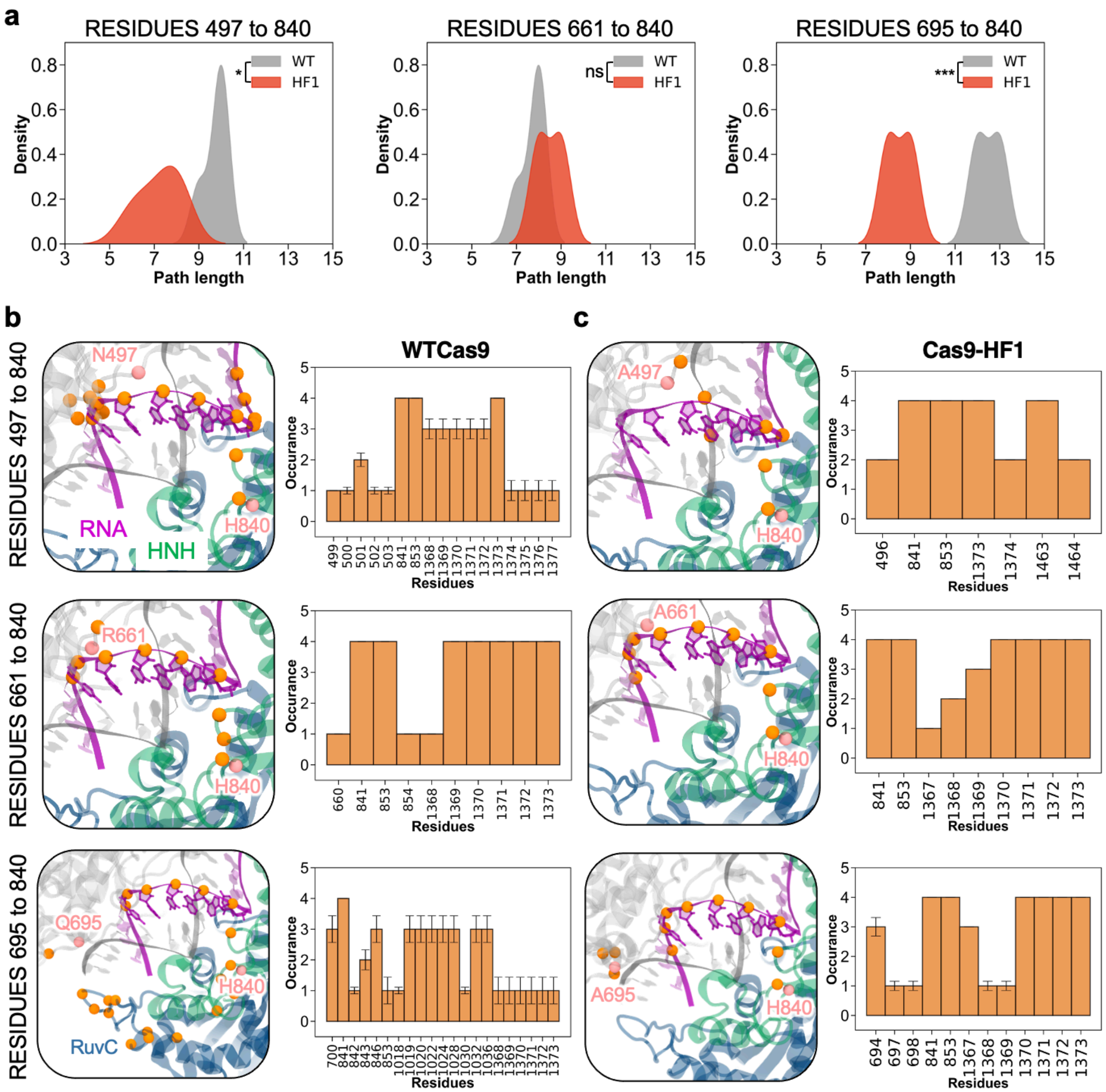


**Supplementary Fig. 14. Analysis of allosteric pathways connecting the Rec3 mutation sites to the HNH catalytic core in the WT and Cas9-HF1. a.** Kernel density estimation of pathlengths in terms of number of edges. Pathways are computed connecting the Rec3 mutation sites (residues 497, 661, and 695) to the HNH catalytic residue H840 and compared between HF1 (red) and WT (grey) Cas9. The statistical significance among the distributions was calculated using a two-tailed unpaired t-test (P-value reference: not significant, ns $P > 0.05$, * $P \leq 0.05$, ** $P \leq 0.005$, *** $P \leq 0.0005$). **b-c.** Occurrence of residues in the optimal and top five suboptimal pathways communicating the Rec3 mutation sites to the HNH catalytic site in the WT (**b**) and HF1 (**c**) Cas9. Error bars correspond to the standard error of the mean computed over three sample pools obtained from ~16μs of aggregated sampling for each of the systems. Signalling pathways are also shown on the three-dimensional structure using spheres of different colours: pink (source and sink) and orange (path nodes).

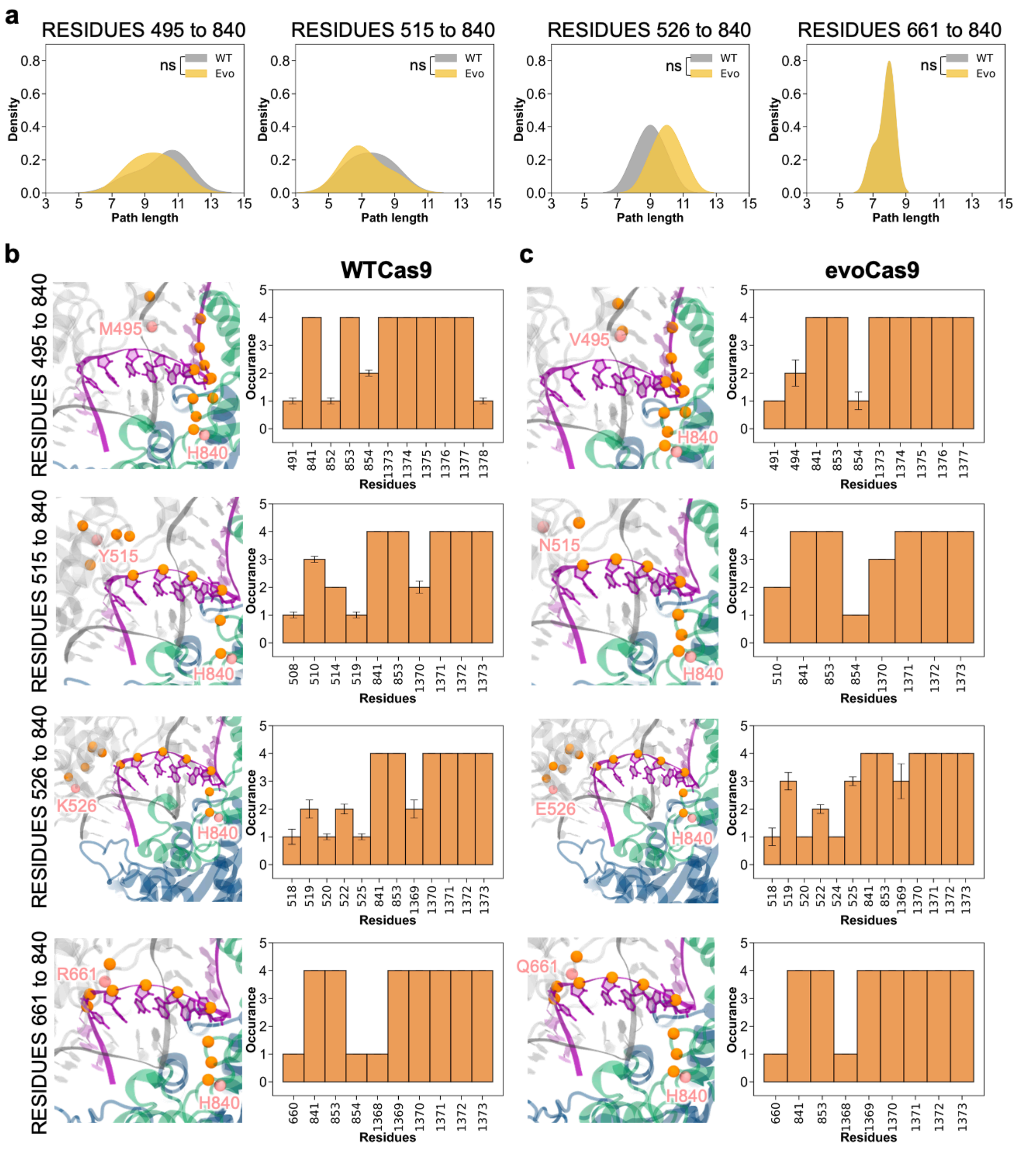


**Supplementary Fig. 15. Analysis of allosteric pathways connecting the Rec3 mutation sites to the HNH catalytic core in the WT and evoCas9. a**. Kernel density estimation of pathlengths in terms of number edges. Pathways are computed connecting the Rec3 mutation sites (residues 495, 515, 526, and 661) to the HNH catalytic residue H840 and compared between Evo (yellow) and WT (grey) Cas9. The statistical significance among the distributions was calculated using a two-tailed unpaired t-test (P-value reference: not significant, ns $P > 0.05$, * $P \leq 0.05$, ** $P \leq 0.005$, *** $P \leq 0.0005$). **b-c.** Occurrence of residues in the optimal and top five suboptimal pathways communicating the Rec3 mutation sites to the HNH catalytic site in the WT (**b**) and Evo (**c**) Cas9. Error bars correspond to the standard error of the mean computed over three sample pools obtained from ~16μs of aggregated sampling for each of the systems. Signalling pathways are also shown on the three-dimensional structure using spheres of different colours: pink (source and sink) and orange (path nodes).

### Supplementary Tables

**Table S1.** Conformational exchange parameters for residues in WT Rec3 via CPMG relaxation dispersion experiments. Data were fit to two-state exchange models as described in the *Materials and Methods* section of the main text.

| WT Rec3 | | | | |
|---|---|---|---|---|
| **Residue** | **Model** | $R_2$ ($s^{-1}$) | $R_{ex}$ ($s^{-1}$) | $k_{ex}$ ($s^{-1}$) |
| 498 | 3 | 31.5 ± 0.01 | 1.8 | 486.9 ± 28.2 |
| 506 | 3 | 29.1 ± 0.07 | 4.6 | 295.0 ± 46.2 |
| 512 | 3 | 33.8 ± 0.07 | 4.4 | 185.4 ± 80.0 |
| 514 | 2 | 39.7 ± 0.20 | 7.6 | 500.0 ± 121 |
| 520 | 2 | 39.0 ± 0.03 | 6.7 | 500.0 ± 12.5 |
| 527 | 3 | 41.4 ± 0.15 | 6.8 | 317.9 ± 123.6 |
| 541 | 3 | 34.9 ± 0.03 | 3.8 | 84.9 ± 19.9 |
| 561 | 2 | 33.1 ± 0.06 | 3.9 | 500.0 ± 50.9 |
| 578 | 2 | 34.7 ± 0.07 | 4.2 | 330.3 ± 35.3 |
| 583 | 2 | 33.8 ± 0.05 | 1.5 | 906.4 ± 170.6 |
| 592 | 2 | 30.6 ± 0.29 | 6.7 | 2215.8 ± 376.3 |
| 596 | 2 | 36.3 ± 0.04 | 4.1 | 851.2 ± 41.9 |
| 597 | 3 | 37.2 ± 0.04 | 4.2 | 181.6 ± 72.5 |
| 625 | 2 | 34.8 ± 0.14 | 4.3 | 500.0 ± 197.7 |
| 626 | 2 | 20.2 ± 0.17 | 3.5 | 1328.1 ± 354.1 |
| 636 | 3 | 33.4 ± 0.01 | 3.1 | 273.9 ± 35.6 |
| 640 | 2 | 33.8 ± 0.10 | 4.4 | 862.3 ± 153.9 |
| 651 | 2 | 35.3 ± 0.10 | 7.2 | 500.0 ± 91.3 |
| 652 | 2 | 33.9 ± 0.08 | 4.8 | 500.0 ± 52.1 |
| 653 | 2 | 33.6 ± 0.08 | 5.2 | 895.4 ± 92.7 |
| 655 | 2 | 28.4 ± 0.07 | 4.8 | 2102.6 ± 109.7 |
| 656 | 2 | 31.6 ± 0.07 | 6.0 | 684.8 ± 62.1 |
| 674 | 3 | 31.6 ± 0.03 | 4.4 | 136.6 ± 52.8 |
| 677 | 3 | 36.6 ± 0.01 | 3.4 | 39.2 ± 8.9 |
| 678 | 3 | 32.9 ± 0.16 | 5.7 | 148.3 ± 80.1 |
| 693 | 2 | 36.2 ± 0.07 | 3.3 | 500.0 ± 75.5 |
| 698 | 2 | 35.8 ± 0.09 | 3.1 | 701.2 ± 107.9 |
| 702 | 2 | 32.3 ± 0.10 | 4.5 | 519.5 ± 95.0 |
| 704 | 2 | 39.7 ± 0.16 | 2.6 | 1194.8 ± 401.3 |
| 708 | 2 | 37.6 ± 0.12 | 5.1 | 500.0 ± 95.5 |
| 710 | 3 | 34.7 ± 0.06 | 2.6 | 328.0 ± 70.4 |
| 712 | 2 | 32.3 ± 0.03 | 2.1 | 122.4 ± 33.1 |

**Table S2.** Conformational exchange parameters for residues in HF1 Rec3 via CPMG relaxation dispersion experiments. Data were fit to two-state exchange models as described in the *Materials and Methods* section of the main text.

| HF1 Rec3 | | | | |
|---|---|---|---|---|
| **Residue** | **Model** | $R_2$ ($s^{-1}$) | $R_{ex}$ ($s^{-1}$) | $k_{ex}$ ($s^{-1}$) |
| 506 | 2 | 26.3 ± 0.05 | 2.9 | 944.5 ± 80.5 |
| 561 | 3 | 29.1 ± 0.06 | 3.1 | 511.8 ± 174.3 |
| 581 | 2 | 27.5 ± 0.06 | 4.4 | 1220.2 ± 105.2 |
| 584 | 3 | 33.0 ± 0.05 | 8.9 | 51.3 ± 24.8 |
| 589 | 2 | 28.6 ± 0.20 | 8.4 | 500.0 ± 84.3 |
| 590 | 2 | 27.1 ± 0.06 | 5 | 500.0 ± 69.1 |
| 626 | 2 | 19.9 ± 0.12 | 3.4 | 344.4 ± 119.2 |
| 628 | 2 | 35.4 ± 0.23 | 3.1 | 1380.2 ± 444.4 |
| 640 | 2 | 30.2 ± 0.04 | 3.9 | 775.6 ± 62.8 |
| 651 | 2 | 31.6 ± 0.06 | 3.8 | 500.0 ± 71.5 |
| 652 | 3 | 30.3 ± 0.08 | 6.1 | 268.3 ± 36.2 |
| 653 | 2 | 30.8 ± 0.14 | 2.8 | 1023.1 ± 207.4 |
| 655 | 2 | 27.6 ± 0.11 | 3.9 | 1545.9 ± 233.3 |
| 656 | 2 | 28.4 ± 0.32 | 5.3 | 940.6 ± 491.3 |
| 659 | 2 | 38.4 ± 0.26 | 6.5 | 500.0 ± 234.0 |
| 662 | 2 | 29.4 ± 0.18 | 4.2 | 610.8 ± 241.7 |
| 669 | 3 | 29.2 ± 0.04 | 2.5 | 826.2 ± 182.1 |
| 670 | 2 | 28.8 ± 0.01 | 1.9 | 500.0 ± 9.2 |
| 674 | 2 | 27.9 ± 0.08 | 3.2 | 1018.8 ± 172.6 |
| 677 | 2 | 32.7 ± 0.06 | 3.3 | 500.0 ± 63.7 |
| 687 | 3 | 45.4 ± 0.07 | 23.4 | 200.5 ± 25.2 |
| 704 | 2 | 36.1 ± 0.01 | 4.9 | 500.0 ± 11.2 |
| 710 | 2 | 30.7 ± 0.09 | 3.4 | 329.9 ± 48.5 |
| 711 | 2 | 30.5 ± 0.18 | 3.5 | 1682.8 ± 292.6 |
| 713 | 2 | 11.1 ± 0.05 | 2.3 | 1331.5 ± 176.4 |

**Table S3.** Conformational exchange parameters for residues in Hypa Rec3 via CPMG relaxation dispersion experiments. Data were fit to two-state exchange models as described in the *Materials and Methods* section of the main text.

| Hypa Rec3 | | | | |
|---|---|---|---|---|
| Residue | Model | $R_2$ ($s^{-1}$) | $R_{ex}$ ($s^{-1}$) | $k_{ex}$ ($s^{-1}$) |
| 498 | 2 | 31.5 ± 0.17 | 5.9 | 624.9 ± 130.7 |
| 504 | 2 | 39.1 ± 0.17 | 6.5 | 1023.6 ± 129.0 |
| 506 | 2 | 28.2 ± 0.11 | 10.1 | 823.7 ± 60.1 |
| 507 | 2 | 34.1 ± 0.42 | 4.3 | 2775.9 ± 521.9 |
| 508 | 2 | 37.2 ± 0.33 | 5.6 | 1586.6 ± 450.8 |
| 510 | 2 | 35.6 ± 0.34 | 16.1 | 500.0 ± 94.0 |
| 511 | 2 | 35.8 ± 0.28 | 8.8 | 500.0 ± 104.9 |
| 514 | 2 | 37.4 ± 0.22 | 5.2 | 514.7 ± 86.5 |
| 516 | 2 | 38.3 ± 0.08 | 4.0 | 500.0 ± 74.1 |
| 518 | 2 | 38.8 ± 0.22 | 7.7 | 209.9 ± 49.8 |
| 520 | 2 | 36.9 ± 0.26 | 7.4 | 592.4 ± 218.8 |
| 521 | 3 | 39.3 ± 0.07 | 5.4 | 307.3 ± 39.7 |
| 528 | 2 | 29.9 ± 0.05 | 8.8 | 754.7 ± 22.5 |
| 540 | 2 | 32.5 ± 0.12 | 4.5 | 551.6 ± 94.3 |
| 541 | 2 | 35.8 ± 0.12 | 3.8 | 500.0 ± 197.9 |
| 550 | 2 | 34.2 ± 0.09 | 2.0 | 500.0 ± 180.4 |
| 551 | 2 | 33.6 ± 0.10 | 5.2 | 500.0 ± 42.9 |
| 554 | 3 | 34.3 ± 0.06 | 4.0 | 239.2 ± 37.2 |
| 556 | 3 | 32.3 ± 0.16 | 7.4 | 405.5 ± 73.6 |
| 583 | 3 | 31.3 ± 0.03 | 4.4 | 234.1 ± 78.6 |
| 590 | 2 | 29.1 ± 0.09 | 5.3 | 324.7 ± 43.0 |
| 591 | 2 | 33.1 ± 0.32 | 10.9 | 500.0 ± 111.1 |
| 592 | 3 | 29.0 ± 0.06 | 6.7 | 16.5 ± 7.5 |
| 596 | 3 | 36.9 ± 0.07 | 5.9 | 170.6 ± 17.4 |
| 597 | 2 | 36.2 ± 0.09 | 4.1 | 501.8 ± 40.4 |
| 607 | 2 | 33.1 ± 0.06 | 1.7 | 500.0 ± 179.6 |
| 609 | 2 | 31.9 ± 0.08 | 5.0 | 500.0 ± 64.1 |
| 615 | 2 | 32.4 ± 0.14 | 5.2 | 500.0 ± 160.8 |
| 624 | 2 | 36.9 ± 0.16 | 2.9 | 603.9 ± 146.7 |
| 625 | 2 | 32.4 ± 0.26 | 2.8 | 500.0 ± 383.6 |
| 626 | 2 | 24.1 ± 0.06 | 8.9 | 778.7 ± 32.0 |
| 640 | 2 | 31.6 ± 0.11 | 5.2 | 698.7 ± 113.6 |
| 647 | 2 | 30.4 ± 0.07 | 3.4 | 286.6 ± 54.7 |
| 652 | 2 | 32.5 ± 0.23 | 5.0 | 1107.7 ± 276.8 |
| 653 | 2 | 31.9 ± 0.16 | 4.7 | 905.8 ± 203.5 |
| 655 | 2 | 27.9 ± 0.05 | 5.9 | 1027.1 ± 44.4 |
| 666 | 2 | 36.3 ± 0.40 | 9.2 | 965.9 ± 239.1 |
| 669 | 2 | 32.2 ± 0.09 | 7.8 | 725.2 ± 50.3 |
| 674 | 2 | 31.1 ± 0.09 | 6.7 | 1475.0 ± 90.7 |
| 676 | 2 | 34.7 ± 0.93 | 5.4 | 2023.7 ± 597.4 |
| 677 | 3 | 35.1 ± 0.14 | 6.7 | 288.6 ± 113.3 |

| | | | | |
|---|---|---|---|---|
| 678 | 2 | 30.5 ± 0.21 | 8.5 | 1885.3 ± 158.4 |
| 679 | 2 | 38.5 ± 0.14 | 4.7 | 500.0 ± 37.7 |
| 680 | 2 | 38.1 ± 0.16 | 5.5 | 541.7 ± 83.6 |
| 681 | 2 | 36.4 ± 0.07 | 5.6 | 597.2 ± 32.1 |
| 682 | 2 | 32.9 ± 1.39 | 8.4 | 3744.5 ± 803.0 |
| 704 | 2 | 34.3 ± 2.47 | 6.7 | 4704.3 ± 1267.8 |
| 706 | 2 | 33.5 ± 0.28 | 7.3 | 1263.9 ± 258.2 |
| 710 | 2 | 33.9 ± 0.17 | 3.5 | 1422.0 ± 331.8 |
| 711 | 2 | 32.9 ± 2.10 | 8.7 | 2994.6 ± 1182.2 |
| 712 | 2 | 29.7 ± 1.89 | 8.7 | 3642.7 ± 963.5 |
| 713 | 3 | 13.0 ± 0.01 | 8.2 | 24.1 ± 1.48 |

**Table S4.** Conformational exchange parameters for residues in WT Rec3 via CPMG relaxation dispersion experiments. Data were fit to two-state exchange models as described in the *Materials and Methods* section of the main text.

| Evo Rec3 | | | | |
|---|---|---|---|---|
| **Residue** | **Model** | **$R_2$ (s$^{-1}$)** | **$R_{ex}$ (s$^{-1}$)** | **$k_{ex}$ (s$^{-1}$)** |
| 506 | 3 | 26.7 ± 0.03 | 5.4 | 12.8 ± 1.11 |
| 527 | 2 | 26.9 ± 0.43 | 4.1 | 3311.5 ± 606.7 |
| 531 | 3 | 27.6 ± 0.02 | 2.8 | 155.5 ± 62.7 |
| 561 | 2 | 29.7 ± 0.07 | 3.7 | 524.6 ± 66.1 |
| 576 | 2 | 34.6 ± 0.33 | 3.6 | 2073.5 ± 822.3 |
| 577 | 2 | 29.4 ± 0.04 | 4.0 | 500.0 ± 26.3 |
| 584 | 3 | 28.1 ± 0.05 | 9.5 | 73.7 ± 21.0 |
| 602 | 2 | 33.6 ± 0.02 | 4.5 | 500.0 ± 18.2 |
| 618 | 2 | 31.4 ± 0.05 | 3.6 | 500.0 ± 64.3 |
| 619 | 2 | 31.6 ± 0.05 | 3.0 | 500.0 ± 155.3 |
| 626 | 2 | 18.7 ± 0.03 | 4.0 | 367.3 ± 31.4 |
| 629 | 2 | 48.6 ± 1.15 | 13.8 | 1126.3 ± 678.6 |
| 640 | 2 | 31.0 ± 0.06 | 4.1 | 937.1 ± 149.6 |
| 650 | 2 | 28.8 ± 0.05 | 2.6 | 500.0 ± 205.1 |
| 652 | 2 | 31.2 ± 0.09 | 5.9 | 502.9 ± 64.0 |
| 653 | 2 | 31.6 ± 0.02 | 3.1 | 867.8 ± 29.7 |
| 654 | 3 | 27.8 ± 0.03 | 6.5 | 16.8 ± 2.87 |
| 656 | 2 | 28.6 ± 0.12 | 5.4 | 1759.7 ± 204.3 |
| 657 | 3 | 26.9 ± 0.03 | 5.3 | 21.1 ± 0.4 |
| 665 | 2 | 35.9 ± 0.04 | 2.2 | 500.0 ± 116.5 |
| 666 | 2 | 35.4 ± 0.08 | 2.5 | 1022.7 ± 163.8 |
| 667 | 2 | 34.3 ± 0.30 | 6.4 | 896.0 ± 427.0 |
| 668 | 3 | 34.7 ± 0.06 | 3.2 | 523.0 ± 156.7 |
| 669 | 3 | 29.5 ± 0.10 | 5.2 | 987.1 ± 158.7 |
| 671 | 2 | 27.6 ± 0.01 | 3.7 | 899.4 ± 15.9 |
| 674 | 3 | 28.6 ± 0.06 | 4.1 | 447.5 ± 64.5 |
| 675 | 2 | 33.7 ± 0.09 | 7.2 | 263.6 ± 27.3 |
| 676 | 2 | 32.4 ± 0.87 | 4.2 | 3262.5 ± 1111.3 |
| 677 | 3 | 34.7 ± 0.06 | 4.3 | 225.9 ± 33.9 |
| 678 | 2 | 29.3 ± 0.05 | 5.0 | 515.1 ± 37.7 |
| 679 | 3 | 37.8 ± 0.20 | 7.7 | 283.3 ± 99.8 |
| 702 | 2 | 29.2 ± 0.01 | 4.8 | 500.0 ± 5.4 |
| 708 | 2 | 33.8 ± 0.20 | 5.5 | 1700.7 ± 283.0 |
| 710 | 2 | 31.2 ± 0.03 | 3.2 | 267.0 ± 20.3 |
| 711 | 2 | 30.9 ± 0.07 | 3.2 | 2551.6 ± 113.5 |
| 712 | 2 | 29.2 ± 0.07 | 2.9 | 1252.9 ± 180.8 |

### Supplementary References

1. Lange, O. F. & Grubmüller, H. Generalized correlation for biomolecular dynamics. *Proteins Struct. Funct. Genet.* **62**, 1053–1061 (2006).

2. Melo, M. C. R., Bernardi, R. C., de la Fuente-Nunez, C. & Luthey-Schulten, Z. Generalized correlation-based dynamical network analysis: a new high-performance approach for identifying allosteric communications in molecular dynamics trajectories. *J. Chem. Phys.* **153**, 134104 (2020).
